# A multidimensional Pan-cancer analysis of CDKN1A identifies CDKN1A as an Immunological and Prognostic Biomarker

**DOI:** 10.1101/2024.09.03.610958

**Authors:** Wenyang Zhang, Qinglong Ma, Wenrun Li, Honghui Zhao, Linghui Zhong, Yinan Xiao, Yaru Ren, Kaixin Yang, Yonghong Li, Lei Shi

## Abstract

CDKN1A/p21 is well recognized for its role in cell cycle regulation and genomic stability. However, its functions in the Tumor microenvironment (TME) and tumor immunity are not yet fully understood. Hereby, we explored CDKN1A expression and immunological/prognostic values via various databases and analytical methods including cBioPortal, Kaplan-Meier, UCSCXenaShiny, TIMER, Single-cell RNA sequencing (scRNA-seq) analysis, etc. In addition, we explored different approaches including CCK8, EdU, Colony formation, Drug sensitivity and Annixin-V assay to explore the influence of p21 in proliferative capacity in cancer cells. We found that CDKN1A is lowly expressed in BLCA, BRCA, COAD, KICH, LUAD, LUSC, PRAD, READ and STAD compared to normal samples, whereas it is highly expressed in CHOL, HNSC, KIRC, KIRP and THCA compared to normal cohorts. CDKN1A expression is significantly correlated with overall survival, disease-specific survival, disease-free survival and progression-free interval different cancer types. Additionally, CDKN1A is associated with CD4+ T cell, CD8+ T cell, Neutrophil, Macrophage and Myeloid dendritic cell infiltration in diverse cancer types. Functional experiments reveal that p21 overexpression leads to a significant reduction in proliferative capacity, facilitates cell apoptosis and senescence in multiple cancer cell lines. In contrast, silenced p21 facilitates cell growth and wound closure, prevent cell senescence in different cancer cell lines. In conclusion, our findings suggest that CDKN1A may serve as a valuable prognostic and immunotherapeutic marker in diverse cancer.

## 1. Introduction

Cancer has become a major global public health issue [1]. According to the World Health Organization (WHO) statistics for 2022, there were approximately 10 million cancer-related deaths, and the number of cancer patients is constantly increasing, profoundly impacting on socioeconomic development, population life expectancy, and quality of life [2]. Tumorigenesis is a dynamic process involving multi-level responses to biological activities, characterized by regulation of genomic instability, epigenetic modifications and irregular cell signaling [3]. The gain-of-function mutations in proto-oncogenes, which strengthen cell proliferation, division and metastasis, along with loss-of-function in tumor suppressor genes, which inhibit tumor development and induce cell cycle arrest, account for the majority of genetic alterations in cancer [4, 5]. Tumor development can arise from lesions affecting the expression or activities of oncogenes, or through the silencing of tumor suppressor genes [6]. Current cancer treatment methods include surgery, chemotherapy, radiation therapy and targeted therapy. First-line options often involve adoptive T cell transfer, agonistic antibodies, cancer vaccines and immunotherapies utilizing immune checkpoint inhibitors [7–9]. Therefore, searching for new targets for immunotherapy through cellular-based research and *in silico* analysis is crucial and indispensable.

Loss of cell cycle control is a hallmark of tumorigenesis [10]. The cell-dependent kinase inhibitor 1A (CDKN1A) gene, which codes for the cyclin-dependent kinase inhibitor p21/WAF1/CIP1/CDKN1A, is the first known target of p53 that inhibits cyclin-CDK complexes largely in G1 arrest [11]. Since its discovered in 1993, subsequent efforts have elucidated the essential functions in cancer phenotype and physiological differentiation. CDKN1A/p21 is responsible for different molecular biological functions such as differentiation, cell migration/invasion, apoptosis, DNA repair and stem cell reprogramming [12]. Emerging studies have demonstrated that p21 can serve as either a tumor suppressor or an oncogene, largely depending on the TME, cell panel, subcellular localization and histone modification [13]. Nevertheless, challenges persist in fully elucidating underlying mechanism and the functions in p53-dependent and p53-independent complex, particularly in tumor immunity.

Our study represents the first comprehensive pan-cancer analysis of CDKN1A, leveraging data from the TCGA project and GEO databases. Utilizing multiple online resources including TCGA, cBioPortal, The Human Protein Atlas (HPA), The UALCAN database, Tumor Immune Estimation Resource (TIMER), Kaplan-Meier Plotter, CancerSEA and others, we investigated the diverse roles of CDKN1A in tumor prevalence and immunological response. First, we examined gene mutations and expression value of CDKN1A/p21 in various cancer types and its association with clinical outcomes. Subsequently, we evaluated the functional implications of CDKN1A/p21 in tumor mutational burden (TMB), microsatellite instability (MSI), DNA methylation, immune infiltration. Additionally, we investigated the expression patterns and interactions of CDKN1A/p21 with different cell populations via scRNA-seq analysis. Furthermore, we conducted multiple analyses to examine the biological function and pathway enrichment of CDKN1A/p21 using gene set enrichment analysis (GSEA), Gene Ontology (GO) and Kyoto Encyclopedia of Genes and Genomes (KEGG). In summary, our study offers a thorough investigation aiming at deepening our understanding of the prognostic values of CDKN1A/p21 in the tumor microenvironment.

## 2. Materials and methods

### Cell Lines and Culture

The lung cancer cell lines H1299 and A549 were cultured in RPMI-1640 medium, while the Breast cancer cell lines MCF-7 and BT474, as well as the pancreatic cancer cell lines Mia and PDC0034 cells were cultured in Dulbecco’s modified Eagle’s medium in supplemented with 10% Fetal Bovine Serum and 100 units/mL penicillin/streptomycin. All cell lines were purchased from Wuhan Pricella Biotechnology Co., Ltd. and cultured at 37 °C in a humidified environment containing 5% CO2. All cell lines were validated by STR profiling and checked for Mycoplasma through inhouse testing.

### Reagents and Antibodies

Cisplatin (HY-17394), and Trametinib (HY-10999) were purchased from MedChemExpress. Antibodies including p21 (10355-1-AP, 1:2000), GCNF (12712-1-AP, 1:1000), CCND3 (26755-1-AP, 1:1000), GAPDH (60004-1-Ig, 1:100000), α-Tubulin (66031-1-Ig, 1:50000), HRP-conjugated Affinipure Goat Anti-Mouse IgG(H+L) (SA00001-1, 1:10000), HRP-conjugated Affinipure Goat Anti-Rabbit IgG(H+L) (SA00001-2, 1:10000) were purchased from Proteintech Group Inc. CD137 (A22167, 1:1000) was purchased from Company ABclonal Inc.

### siRNA and Plasmid Transfection

siRNAs were designed and purchased from Guangzhou RiboBio Co., Ltd. siRNA (50 nm) was transfected using Hiperfect (Qiagen) for 48 hours according to the manufacturer’s protocol. Plasmid was transiently transfected into cells for 48 hours with lipofectamine 2000 (Mei5bio) reagent according to the manufacturer’s protocol.

### RNA extraction and RT-qPCR

Total RNA was isolated using the M5 Total RNA Extraction Reagent (Mei5bio). cDNA was generated from 1 µg of total RNA per sample using M5 Sprint qPCR RT kit (Mei5bio). Quantitative real-time PCR (RT-qPCR) was performed by using a Light Cycler 480 (Roche) and the Fast SYBR Green Master Mix (Mei5bio). The qPCR primer sequences are shown in table I.

### Protein extraction and Western Blot

Total protein was homogenized in 1 × RIPA buffer (Boster) plus with protease inhibitors and phosphatase inhibitors for 20 minutes, and subsequently centrifuged at 12,000 rpm for 20 minutes at 4 °C. The western blot was performed as previously described [14].

### Annexin-V assay

For apoptosis analysis, H1299, A549, MCF-7, BT474, Mia and PDC0034 cells were transfected with p21 plasmid in 6-well plates for 48 hours. Cells were stained using the Annexin V-FITC/PI Apoptosis kit and assayed according to the instructions (Elabscience). The percentage of apoptotic cells was analyzed using a flow cytometer (BD Accuri C6).

### Colony formation assay

2 × 10^3^ pre-treated cells were seeded in each well of six-well plates. 14 days later, cells were subsequently washed with PBS and fixed with cold methanol, stained with 0.05% crystal violet (BKMAM), and counted using the ImageJ software.

### Wound healing assay

To stop cell proliferation, mitomycin C (10 ng/ml) was applied 2 hours before wound generation. After remove the Culture-insert (IBIDI), cells were washed twice with PBS solution to remove mitomycin C and detached cells. The cells were allowed to close the wound for 24-48 hours, and images were captured and evaluated microscopically.

### 5-Ethynyl-2’-deoxyuridine Assay (EDU)

The cells were incubated with fresh prepared EDU buffer (Ribobio) for 2 h. After three washes with PBS, the cells were incubated with 100 μL of 1 × Apollo reaction cocktail for 30 min. Then, the cells were washed three times with 0.5% Triton X-100. The DNA contents were stained with 100 μL of 1 × Hoechst 33 342 (5 μg/mL) for 30 min and visualized under a fluorescence microscope.

### β-Galactosidase Staining

Cell senescence was assessed using β-galactosidase (SA-β-gal) staining kits (Beyotime Biotechnology). Briefly, the cells were washed once with PBS and fixed with β-galactosidase staining fixative solution for 15 min at room temperature. After 3 times with PBS, cells were then incubated with SA-β-gal staining solution at 37 °C overnight, followed with observation under an optical microscope.

### CDKN1A expression in diverse cancer types

The expression level of CDKN1A among various cancer types was examined utilizing the public database UCSC Xena [15], TIMER [16] and UALCAN [16], respectively. Transcriptomic RNA-seq data, DNA methylation data and phenotype data for 33 cancers were obtained from TCGA database. These cancers include Adrenocortical Carcinoma (ACC), Bladder Urothelial Carcinoma (BLCA), Breast Invasive Carcinoma (BRCA), Colon Adenocarcinoma (COAD), Lymphoid Neoplasm Diffuse Large B-cell Lymphoma (DLBC), Esophageal Carcinoma (ESCA), Glioblastoma Multiforme (GBM), Head and Neck Squamous Cell Carcinoma (HNSC), Kidney Chromophobe (KICH), Kidney Renal Clear Cell Carcinoma (KIRC), Kidney Renal Papillary Cell Carcinoma (KIRP), Acute Myeloid Leukemia (LAML), Brain Lower Grade Glioma (LGG), Liver Hepatocellular Carcinoma (LIHC), Lung Adenocarcinoma (LUAD), Lung Squamous Cell Carcinoma (LUSC), Ovarian Serous Cystadenocarcinoma (OV), Pancreatic Adenocarcinoma (PAAD), Prostate Adenocarcinoma (PRAD), Rectum Adenocarcinoma (READ), Skin Cutaneous Melanoma (SKCM), Stomach Adenocarcinoma (STAD), Testicular Germ Cell Tumors (TGCT), Thyroid Carcinoma (THCA), Thymoma (THYM), Uterine Corpus Endometrial Carcinoma (UCEC), Cholangiocarcinoma (CHOL), Cervical Squamous Cell Carcinoma and Endocervical Adenocarcinoma (CESC), Mesothelioma (MESO), Uveal Melanoma (UVM), Pheochromocytoma and Paraganglioma (PCPG), Sarcoma (SARC), Uterine Carcinosarcoma (UCS).

Furthermore, we utilized the HPA database (https://www.proteinatlas.org/) to examine the differential expression of CDKN1A at the protein level [17].

### Mutation analysis of CDKN1A

The pan-cancer data from the cBioPortal database was employed to investigate and compile details of CDKN1A mutations across various malignancies, including mutation frequency and type [18].

### Prognostic analysis of CDKN1A in different cancers

UCSCXenaShiny was utilized to examine the correlation between CDKN1A level and the patient prognosis, including overall survival (OS), disease-specific survival (DSS), disease-free survival (DFS) and progression-free interval (PFI) in 33 cancer types [19]. Patients were divided into CDKN1A-High and CDKN1A-Low expression groups within cancer cohorts. Survival curves were plotted by the "survminer" package. A *p* value < 0.05 was considered significant. The R language "forestplot" package was employed to depict forest plots with log-transformed HR values and 95% CIs.

We also used PrognoScan and Kaplan-Meier Plotter database to analyze the correlation between CDKN1A levels and patient survival time in various cancers. PrognoScan, a publicly accessible database containing a large number of cancer datasets, is used to determine the correlation between gene expression and prognostic values such as OS, DFS, DSS and PFI [20]. The Kaplan-Meier Plotter database, which sources data from GEO, EGA and TCGA, was used to analyze the relationship between gene expression and survival, including OS and relapse-free survival (RFS) rates, in 21 cancer types [21]. The UALCAN database was used to analyze the association of CDKN1A gene expression with different tumor stages in multiple cancer types. In order to assess the diagnostic accuracy of CDKN1A, a ROC curve analysis based on sensitivity and specificity was performed using the "ROCR" software package [22]. A *p* value < 0.05 was considered statistically significant.

### Tumor infiltration analysis

We used bioinformatics analysis tool TIMER to analyze the correlation between CDKN1A expression levels and the infiltration abundance of six immune cell types: B-cells, CD4+ T cells, CD8+ T cells, neutrophils, macrophages and dendritic cells. We also used the UCSCXenaShiny database to evaluate the correlation between CDKN1A and different immune cells.

We obtained data for TMB and MSI analyses from the UCSCXenaShiny database. In addition, the correlations between CDKN1A and different MMR genes (including EPCAM, PMS2, MSH2, MSH6 and MLH1) were explored in 33 tumors through the TIMER database. The relationships between CDKN1A and tumor immune molecules, including immune activating molecules and immune suppressing molecules were collected through the TISIDB dataset [23]. The correlation heatmaps were generated by the R package “pheatmap” [24].

### Methylation analysis

The UALCAN database was used to analyze the methylation levels of the CDKN1A promoter. Beta values were applied to indicate DNA methylation levels, ranging from 0 (unmethylated) to 1 (fully methylated). Different cut-off values of Beta were employed to indicate hypermethylation [Beta: 0.5-0.7] or hypomethylation [Beta: 0.25-0.3] in this study. In addition, we retrieved the methylation data of CDKN1A in pan-cancer through the UCSCXenaShiny database to analyze the methylation level of CDKN1A in pan-cancer.

### Gene-gene and protein-protein interaction analysis

GeneMANIA (http://www.genemania.org) is an interactive and intuitive tool to investigate a series of functions, e.g. protein and genetic interactions, signalling enrichment, gene and gene co-expression and co-localization and protein domain similarity [25]. In this study, we used GeneMANIA to determine the novel genes in the up-or downstream CDKNA1A-relevant pathway.

We also used the PINA database (https://omics.bjcancer.org/pina/) to analyze cancer protein interactions. The screening criteria included tumor type specificity score = 2, correlation coefficients = 0.2 (Spearman), and survival comparison between High 50% and LOW 50%. Cancer Drivers and Drug target genes were also mapped on the PPI networks [26].

### Gene signature analysis

The gene expression files for different cancer types were acquired from the web-based tool Broad GDAC Firehose [27]. The data were divided into two subclasses (CDKN1A Low and CDKN1A High) based on the mean level of CDKN1A. Gene set enrichment analysis (GSEA) was then used to determine signature enrichment in different cancer types [28]. A bubble diagram was generated using the R package ggplot2 to visualize the general biological function in BLCA, COAD, LUAD and LUSC.

We further utilized the CancerSEA database to analyze the correlation of CDKN1A with different functional states in 14 cancers [29].

### Single-cell RNA sequencing (scRNA-seq)

We used the scDVA database (http://panmyeloid.cancer-pku.cn/) to analyze the clustering of CDKN1A in 15 cancer types of 210 patients and its relevant correlation with different immune cell types [30]. Six scRNA-seq samples were converted into Seurat objects using the Seurat R package (version 4.30) [31] and the scater R package (version 1.280) [32]. We also downloaded and processed the scRNA-seq data of LUAD from Ocean Code dataset to examine the association of CDKN1A with different cell populations [33].

### Statistical Analysis

All data are presented as the mean value of ± S.D (n=3) otherwise indicated differently. The significances were calculated by two paired t tests. ∗ p < 0.05, ∗∗ p < 0.01, and ∗∗∗ p < 0.001 were defined as statistically significant. Pearson’s correlation was calculated using the GraphPad Prism package (GraphPad Software Inc.)

## 3. Results

### 3.1 Somatic mutation of CDKN1A in cancers

Copy number variation (CNV), which changes in the number of copies of specific segments among individuals, can contribute to cancer development [34]. To initiate our investigation, we examined the copy number of CDKN1A across various cancers. Through the cBioPortal database, we found that CDKN1A is frequently altered in several cancer types, including bladder cancer, melanoma, lung cancer, hepatobiliary cancer, bone cancer, esophagogastric cancer, breast cancer, soft tissue cancer, colorectal cancer, cervical cancer, and uterine endometrioid cancer (Figure. 1A). Notably, bladder cancer tumor samples present the highest frequency of CDKN1A genetic alterations (>20%), with a significant proportion comprising various mutation types. CDKN1A mutations were detected in approximately 10% of BLCA cohorts, followed by KICH, ASC, LIHC, CESC, SKCM, STAD, ESCA (Figure. 1B). Interestingly, CDKN1A amplification accounts more prevalent than mutations in melanoma, lung cancer and hepatobiliary cancer. Conversely, deep deletions, rather than amplifications, are observed in soft tissue cancer (Figure. 1A). Furthermore, we did not observe a relevance between copy number and gene expression in deletion, diploid, gain and amplification cohorts (Figure. 1C, 1D), indicating that CDKN1A copy number does not influence CDKN1A gene expression. Given the lack of relevant studies, our next focus was to explore the expression and functions of CDKN1A in various cancers.

**Figure 1.**
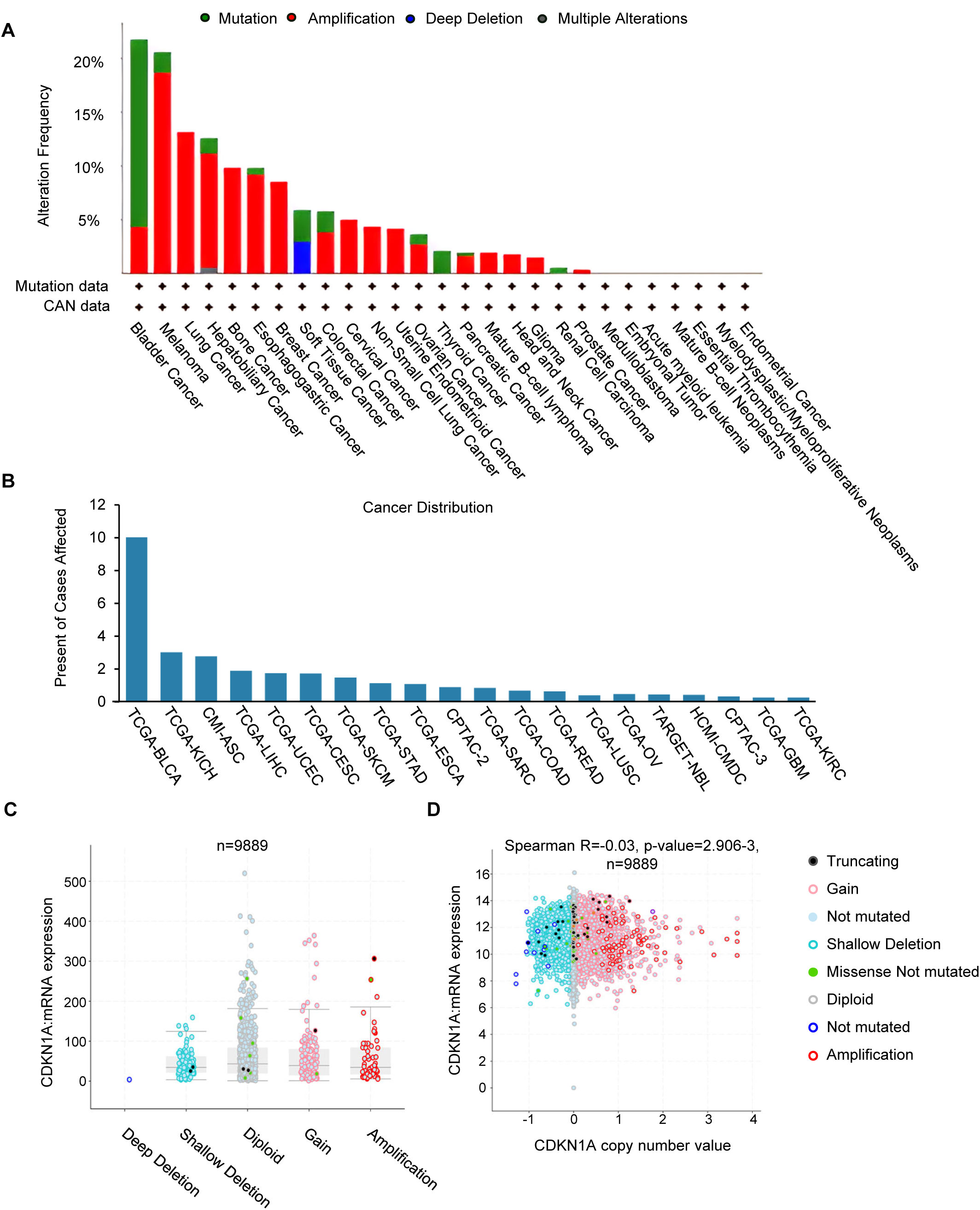
Mutation feature of CDKN1A in pan-cancer from TCGA. (A) The alteration frequency of CDKN1A in pan-cancer datasets through the cBioPortal database. (B) TCGA database showing the mutations of CDKN1A in different cancers. (C) The mRNA level of CDKN1A in tissues with gene copy number deletion, diploid, gain and amplification across all TCGA cancers. (D) The dot plot showing the correlation between CDKN1A copy number and mRNA in pan-cancer.

### 3.2 CDKN1A expression in pan-cancer

To gain better insights into CDKN1A, we first examined its expression in various cancer types. We downloaded the raw TCGA data from UCSC Xena and processed the expression value of CDKN1A among cancers via the R package. CDKN1A is lowly expressed in BLCA, BRCA, COAD, KICH, LUAD, LUSC, PRAD, READ and STAD compared to normal samples, whereas it is highly expressed in CHOL, HNSC, KIRC, KIRP and THCA compared to normal cohorts (Figure. 2A). We found similar results via the TIMER database (Figure. 2B) and UALCAN (Figure. 2C and Table II) databases, respectively.

**Figure 2.**
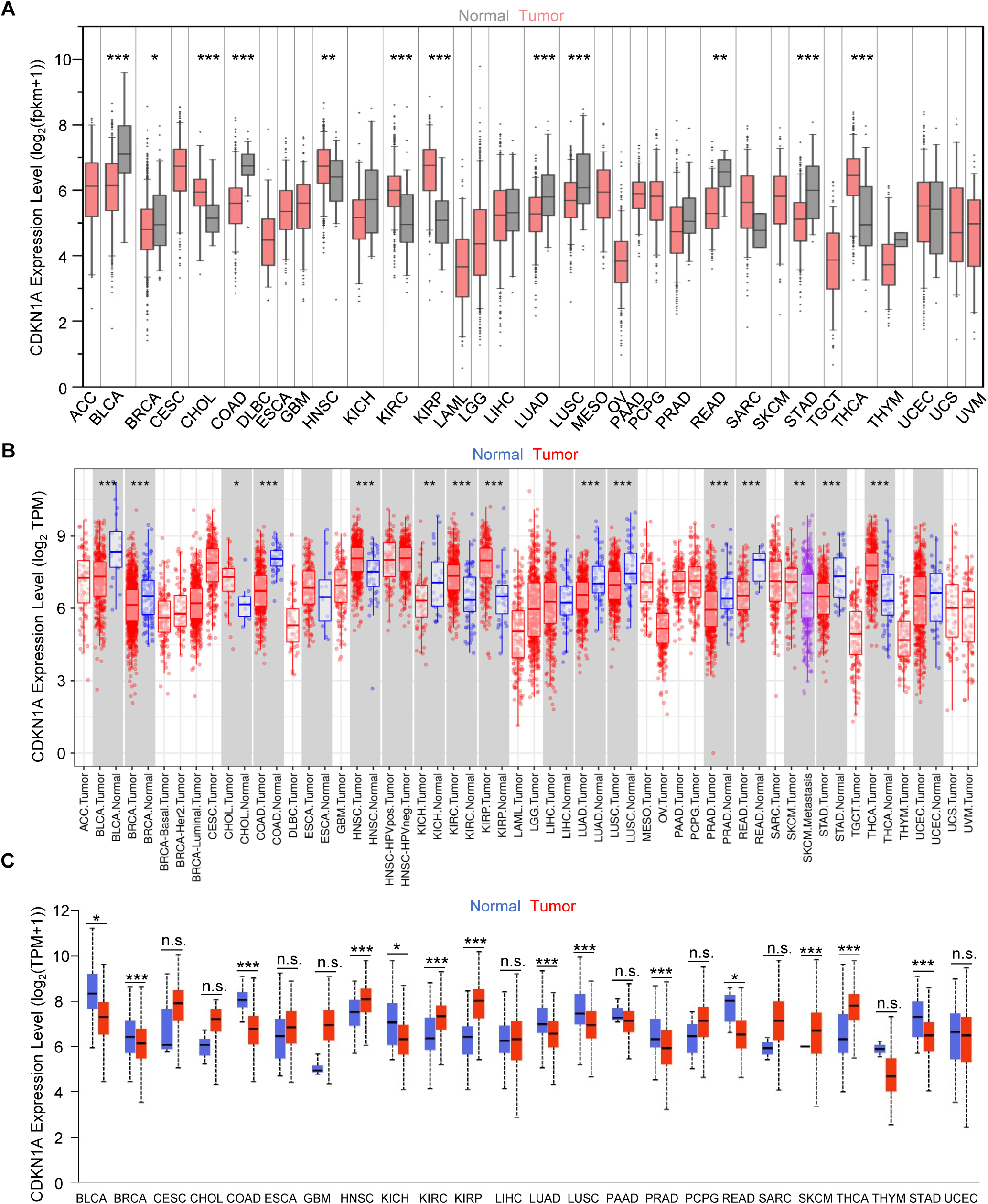
CDKN1A mRNA level in pan-cancer. (A-C) CDKN1A mRNA level from the TCGA database (A), TIMER (B) and UALCAN (C) database. ∗ *p* < 0.05, ∗∗ *p* < 0.01, and ∗∗∗ *p* < 0.001.

In addition, we explored the protein levels of CDKN1A in different cancer types through the UALCAN database. We discovered that CDKN1A protein levels are higher in GBM, HNSC and KIRC compared to normal counterparts (Figure. 3A-C), which matches the mRNA levels in cancers. Immunohistochemical experiments further elucidate the low protein expression of CDKN1A in colon cancer, breast cancer and prostate cancer (Figure. 3E-G), consistent with its mRNA levels in tumor cohorts. Interestingly, although CDKN1A mRNA expression is lower in lung cancer (Figure. 2), its protein levels are higher in tumors, suggesting the existence of post-transcriptional regulation during tumorigenesis (Figure. 3D, 3H).

**Figure 3.**
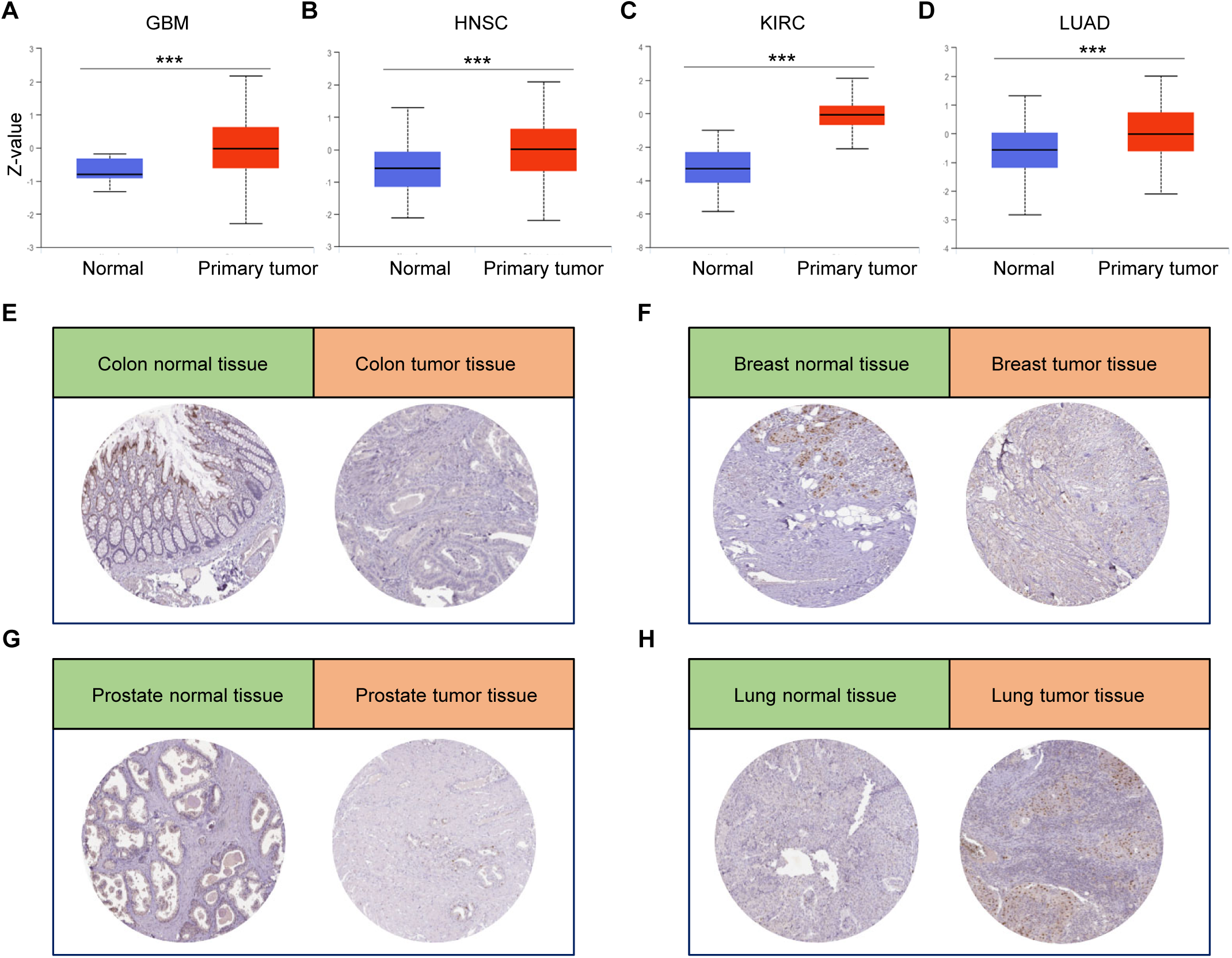
Protein level of p21 in diverse cancers. (A-D) UALCAN dataset indicating the protein level of CDKN1A in GBM (A), HNSC (B), KIRC (C) and LUAD (D). (E-H) Immunohistochemical experiments showing the CDKN1A protein level in colon cancer (E), breast cancer (F), prostate cancer (G) and lung cancer (H). ∗ *p* < 0.05, ∗∗ *p* < 0.01, and ∗∗∗ *p* < 0.001.

### 3.3 Methylation level of CDKN1A

Emerging studies have demonstrated that the dynamic modulation of DNA methylation is an essential molecular mechanism of cancer initiation, maintenance and progression [35]. Therefore, we sought to determine whether the methylation is involved of dysregulation of CDKN1A in pan-cancer. DNA methylation contains two types: one is hypermethylation, which is inversely correlated with gene level and results in the silencing of many genes, and hypomethylation, an early event detected in cancer that promotes gene expression [36]. We found that lower hypomethylation levels of CDKN1A in cancer tissues such as BLCA, CESC, COAD, ESCA, KIRC, KIRP, LIHC, LUAD, LUSC, THCA and UCEC compared to normal tissues, and higher in tissues such as BRCA, CHOL and PRAD (Figure. 4A), which is consistent with the findings obtained from the UALCAN database (Supplementary Figure. 1) and TCGA database (Figure. 4B and Table III).

**Figure 4.**
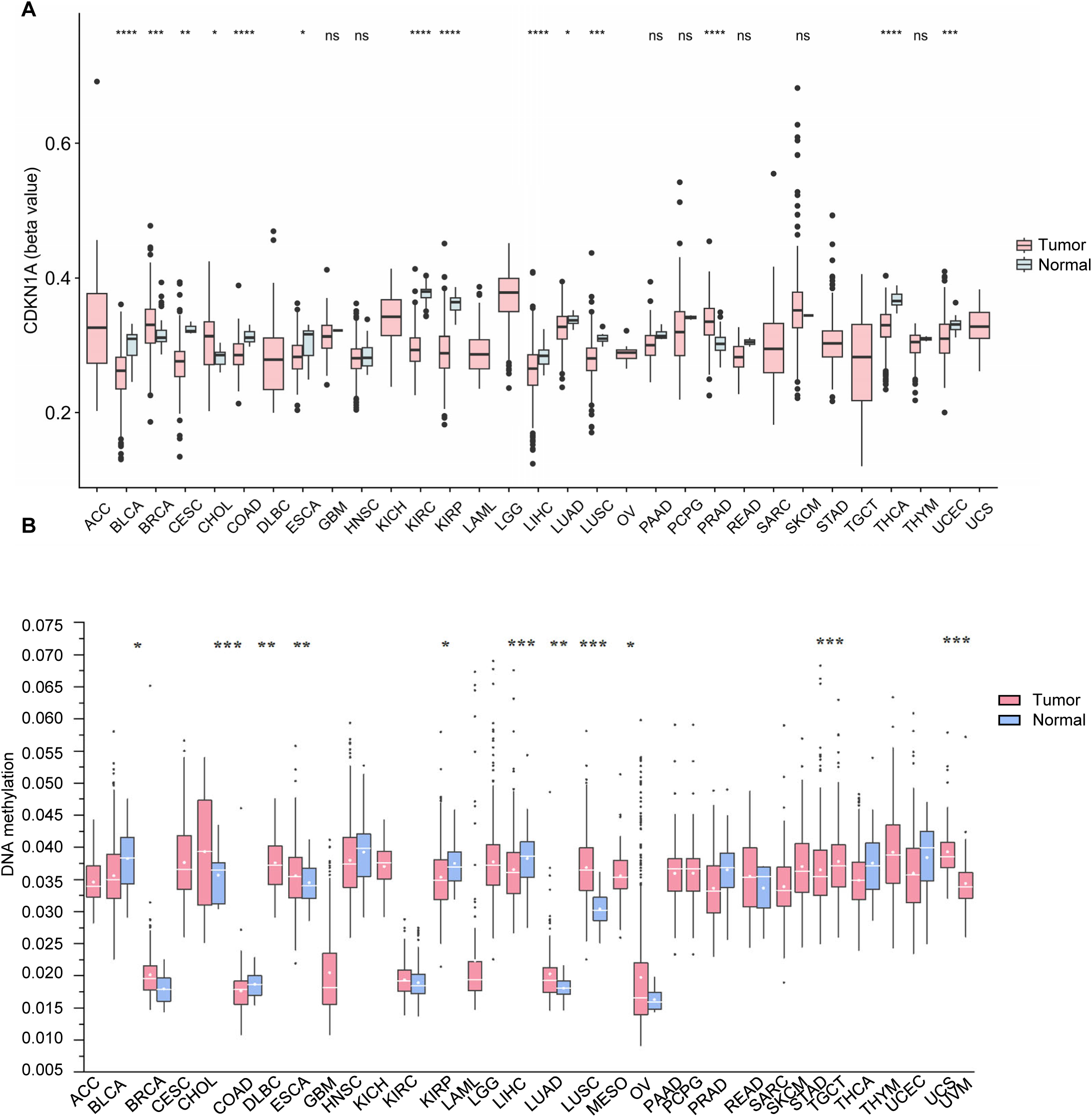
Methylation level of CDKN1A in pan-cancer. (A) CDKN1A methylation of from the UCSCXenaShiny database. CDKN1A has lower levels of hypomethylation in cancer tissues such as BLCA, CESC, COAD, ESCA, KIRC, KIRP, LIHC, LUAD, LUSC, THCA and UCEC, and higher levels of methylation in BRCA, CHOL and PRAD compared to normal samples. (B) CDKN1A has lower methylation levels in LIHC, PRAD, UCEC, BLCA, KIRP, THCA, COAD, and higher methylation levels in LUSC, BRCA, LUAD, and KIRC. ∗ *p* < 0.05, ∗∗ *p* < 0.01, ∗∗∗ *p* < 0.001 and ∗∗∗∗ *p* < 0.0001.

### 3.4 Single cell clustering analysis

Traditional transcriptomics RNA sequencing methods provide average information from a large population of cells, failing to capture the details of cellular heterogeneity. Recent efforts have revealed that scRNA-seq offers notable insights into the heterogeneity of tumor cells, encompassing both the neoplastic compartment and the tumor microenvironment [37]. We therefore carried out the scRNA-seq analysis to examine the CDKN1A relevant clustering and expression in different cell subtypes. We observed the distribution of CDKN1A in 138,161 myeloid cells from 15 cancer types of 210 patients (Figure. 5A, 5B). In general, most of these cell subpopulations are tumor cells (Figure. 5C), and CDKN1A is lowly expressed in tumors compared to normal samples (Figure. 5E). In addition, we found CDKN1A is dramatically distributed in cDC1, cDC2, cDC3, Macro, Mast, Monolike, myeloid and pDC cells (Figure. 5D), with particular high expression in cDC3 cells (Figure. 5F). Interestingly, MYC, a known negative regulator of CDKN1A, is negatively correlated with CDKN1A in different cell types, particularly in cDC3 cell type (Figure. 5G).

**Figure 5.**
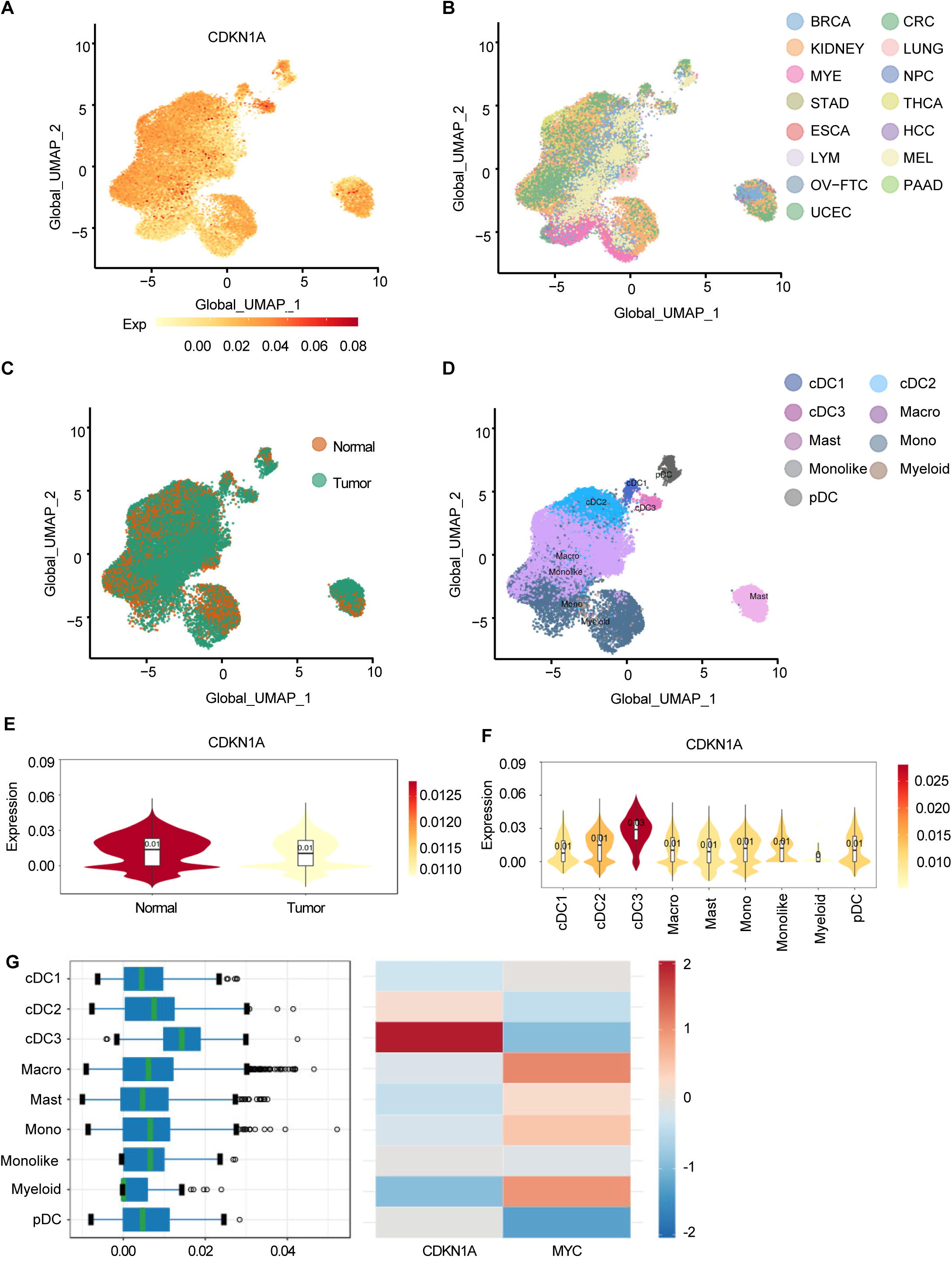
Clustering of CDKN1A in pan-cancer tissues. (A-D) scRNA-seq revealing the CDKN1A gene expression in pan-cancer (A, B), normal/tumor samples (A, C) and cell subpopulations (A, D). (E, F) The expression of CDKN1A in normal/tumor cohort (E) and different cell subtypes (F). (G) The correlation between CDKN1A and MYC in different cell subtypes.

In addition, clustering analysis revealed that the major cellular subpopulation in LUAD tumor cells are AT2 cells and macrophage cells (Supplementary Figure. 2A, 2B) [38]. CDKN1A is highly expressed in normal cells and relatively low in tumor cells (Supplementary Figure. 2A, 2C). Meanwhile, we compared and analyzed the expression of CDKN1A and KRAS in tumor and normal tissues and found that CDKN1A is highly expressed in normal cells, whereas KRAS is highly expressed in tumor samples (Supplementary Figure. 2D). Thus, we deduced that CDKN1A may control distinct cell subpopulations in various malignancies to trigger the onset and progression of cancer.

### 3.5 Gene signature analysis of CDKN1A in pan-cancer

After that, we inquired to understand the CDKN1A-associated gene signatures in various malignancies. We downloaded the TCGA datasets of LUAD, LUSC, BLCA and COAD due to the well-known roles as a tumor suppressor with a low expression in tumor samples (Figure. 2). We divided the datasets into two subcategories (CDKN1A Low and CDKN1A High) based on the mean level of CDKN1A, and performed GSEA analysis using diverse gene sets including Hallmark gene sets, C2 curated gene sets, C5 ontology gene sets, C6 oncogenic signature gene sets and C7 immunological signature gene sets. Our analysis revealed that common signatures, including p53 pathway, Apoptosis, TGF-β signalling, IL6-JAK-STAT3 signalling, KRAS signalling, G2M checkpoint and MYC targets, are enrichened across these four cancer types (Figure. 6A). We next conducted immunoblotting and confirmed that enforced MYC represses p21 in lung cancer and breast cancer (Figure 6B), supporting the fact that p21 may participate in MYC signalling. Additionally, Gene Ontology analysis indicates CDKN1A is associated with DNA repair, Mitotic Spindle, Catalytic Activity in LUSC (Figure. 6C), DNA repair, Cell killing and DNA helicase in COAD (Figure. 6D), Histone binding and Ribosome in BLCA (Figure. 6E), RNA splicing, RNA localization, RNA exportation and Cytokine binding in LUAD (Figure. 6F). Additionally, we discovered that "KRAS", "G2M-CHECKPOINT" and "MYC" are significantly enriched in the datasets of BLCA, COAD, LUAD and LUSC (Supplementary Figure. 3A). Besides, we also found the enrichment of TNF signaling pathway and IL-17 signaling pathway, which are closely related to the immune system in BLCA, p53 signaling, VEGF signaling pathway and IL-17 signaling pathway in LUAD (Supplementary Figure. 3B, 3C). In summary, these gene signature analyses imply the versatile functions of CDKN1A in malignancies.

**Figure 6.**
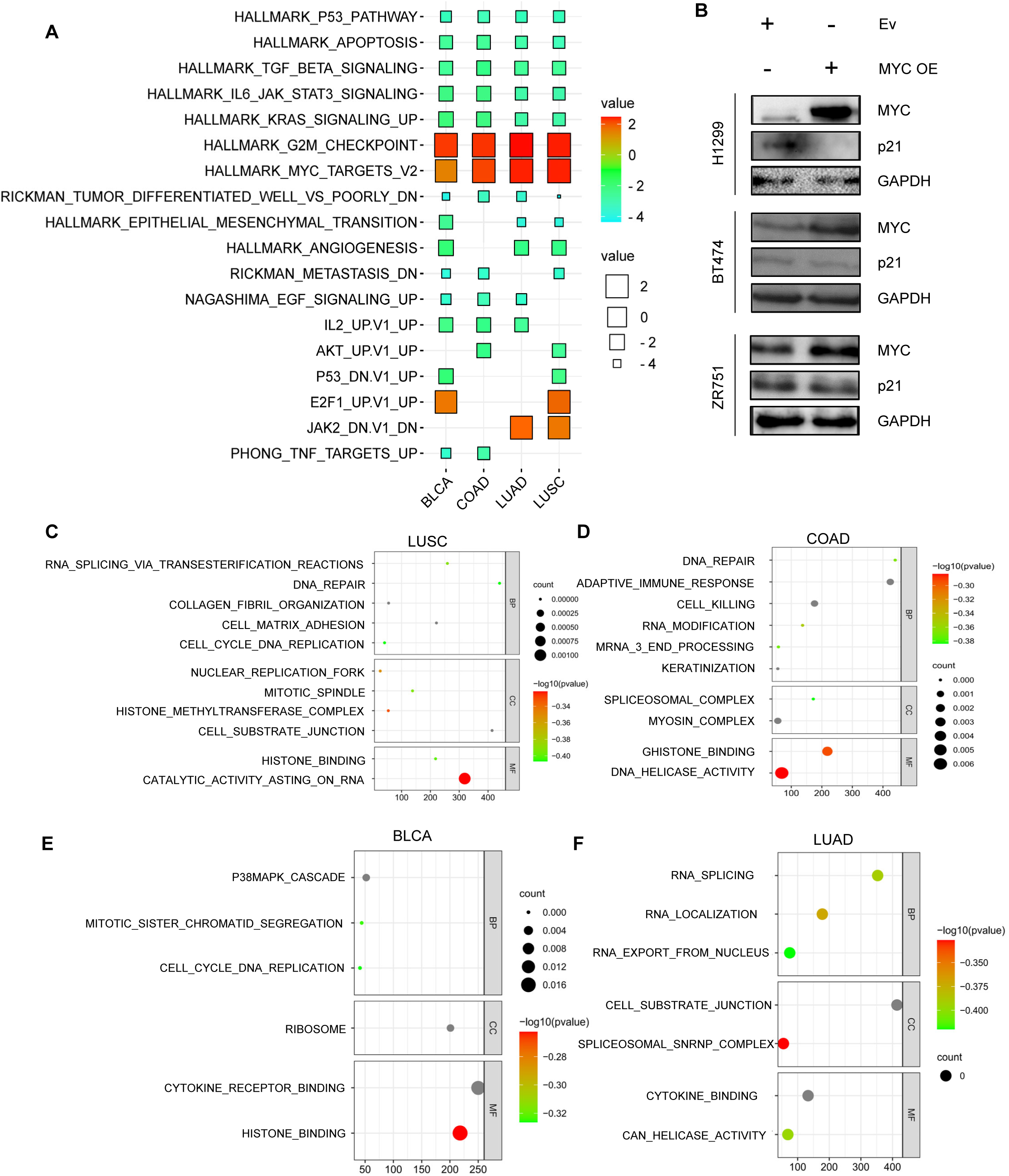
Gene signature analysis of CDKN1A low and CDKN1A high dataset among BLCA, COAD, LUAD and LUSC. (A) Common gene signature among different cancer types. (B) Enforced MYC represses p21 level in H1299, BT474 and ZR751 cells. (C-F) Bubble plots showing GO analysis of CDKN1A Low and CDKN1A High datasets in LUSC (C), COAD (D), BLCA (E), and LUAD (F).

### 3.6 CDKN1A is associated with tumor immunity

The combination of scRNA-seq and GSEA analysis suggests that CDKN1A is associated with TME. To explore this relationship further, we assessed the correlation of CDKN1A with immune cell infiltration using various user-friendly web-based algorithms, including UCSCXenaShiny and the TIMER database. Our analyses show that CDKN1A expression is positively associated with immune infiltrations of CD4 + T cell, CD8 + T cell, Neutrophil, Macrophage and Myeloid dendritic cell infiltration in diverse cancer types (Figure. 7A). TIMER database was used to investigate the enrichment of CDKN1A in cell infiltration with a threshold R > 0.2 and a *p*-value < 0.05. CDKN1A has favorable relationships with CD8 + T cell in UCEC, macrophage in BRCA and dendritic cell in KIRP and UCEC (Supplementary Figure. 4). Furthermore, we found substantial associations between 22 immune cells and CDKN1A in a range of malignancies, including LUAD, LUSC, LAML and THYM (Figure. 7B). We also examined the relationship between CDKN1A and different immunomodulators (immune activators, immunosuppressants) via the TISIDB database. Most immunostimulatory and immunosuppressive genes have strong correlations with CDKN1A expression (Figure. 7C, 7D). In conclusion, our findings highlight a fascinating phenomenon wherein CDKN1A is correlated with genes that both stimulate and repress the immune system in a variety of tumor types. This suggests that CDKN1A may play a fundamental role in shaping the tumor immune microenvironment and could potentially be leveraged in clinical immunotherapy strategies.

**Figure 7.**
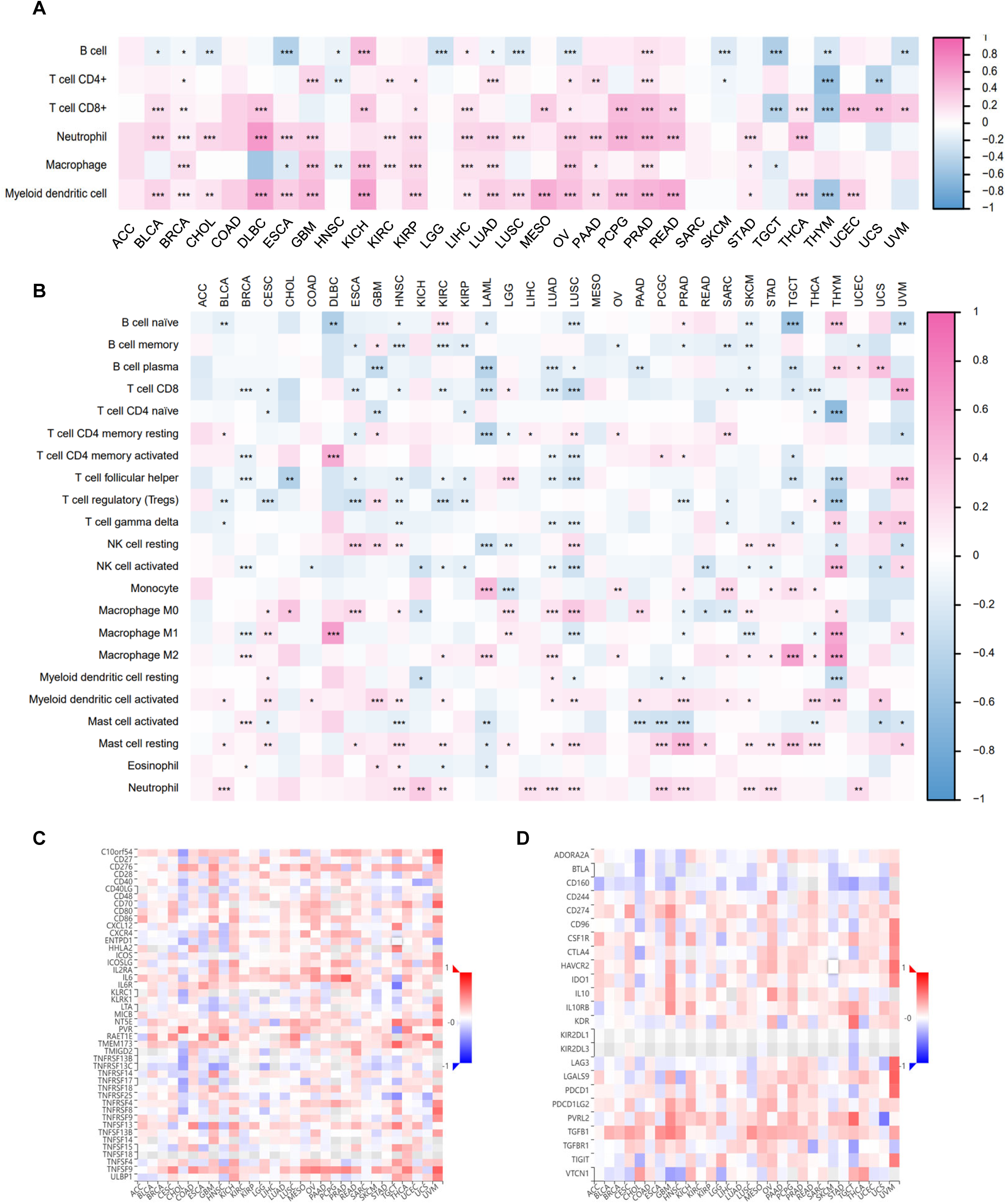
The CDKN1A expression correlated with immune. (A, B) The UCSCXenaShiny database showing the correlation between CDKN1A expression and the infiltration of 6 immune cells (A) and 22 immune cells (B). (C, D) The correlation of CDKN1A expression and immune stimulators (C) and inhibitor (D) in cancers through TISIDB. ∗ *p* < 0.05, ∗∗ *p* < 0.01, and ∗∗∗ *p* < 0.001.

Increasing studies have shown that TMB and MSI are crucial factors in cancer immunology [39, 40]. To explore the relationship between CDKN1A expression and these genomic instability markers, we analyzed TMB and MSI data obtained from the UCSCXenaShiny database. CDKN1A expression is positively correlated with TMB in THYM and UCEC, while negative correlation in GBM, HNSC, LUAD, LUSC, SKCM (Figure. 8A). MSI analysis reveals a negative correlation of CDKN1A with ACC, BLCA, GBM, HNSC, LUAD, PAAD cohorts (Figure. 8B). Considering that mismatch repair (MMR) is a key pathway in maintaining genomic stability and its deficiency leads to MSI-associated tumorigenesis and drug resistance, we evaluated the correlation between CDKN1A expression and five MMR genes [41]. Interestingly, CDKN1A is highly correlated with MMR genes in CHOL, LAML, PAAD, PCPG and LUSC, but not in BLCA, BRCA, DLBC and LGG (Figure. 8C). To validate these findings, we examined the mRNA level of MMR genes after overexpression of p21. We observed that enforced p21 stimulates MSH2, EPCAM and PMS2 expression I in H1299, MCF-7 and BT474 cell (Figure. 8D-8F). Overall, our results suggest that CDKN1A modulates MMR gene expression, potentially contributing to genomic stability in various cancer types.

**Figure 8.**
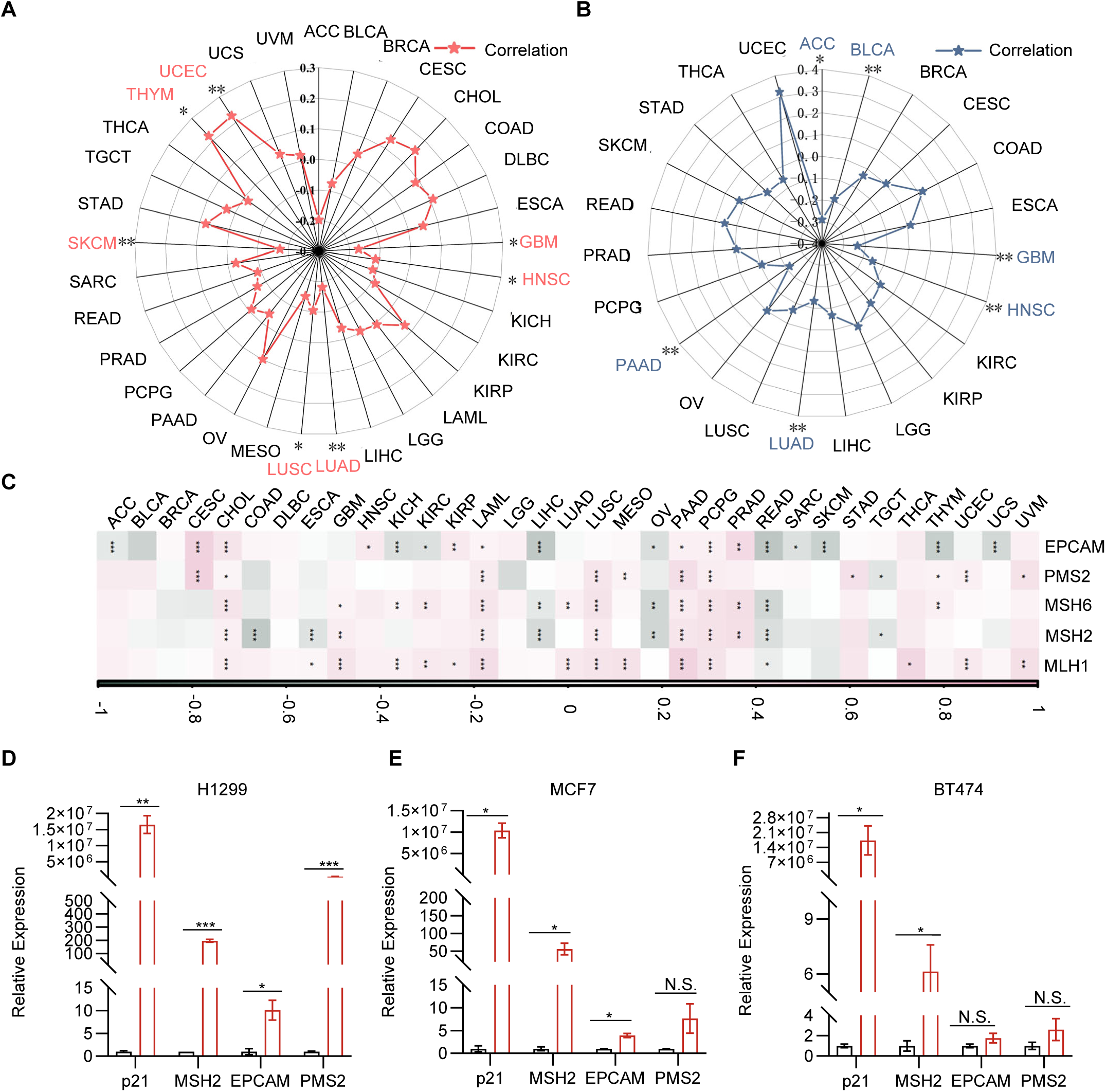
The correlation between CDKN1A with TMB, MSI and MMR. (A) TMB analysis of the correlation between CDKN1A and different cancers. (B) MSI analysis of the correlation between CDKN1A and different cancers. (C) Correlation between CDKN1A expression levels and five MMR genes in diverse cancer. (D-F) qPCR showing MMR gene expression upon p21 overexpression in H1299 (D), MCF-7 (E) and BT474 (F). ∗ *p* < 0.05, ∗∗ *p* < 0.01, and ∗∗∗ *p* < 0.001.

### 3.7 CDKN1A regulates cell proliferation

We further analyzed the correlation between CDKN1A expression and the functional status of 14 cancers using CancerSEA database. In most tumors, CDKN1A is significantly associated with cell-dependent manner activities, hypoxia, inflammation and metastasis (Figure. 9A). In particular, we observed CDKN1A affects apoptosis, cell cycle, differentiation, DNA damage, EMT, inflammation, invasion, metastasis in cancers (Figure. 9B, 9C).

**Figure 9.**
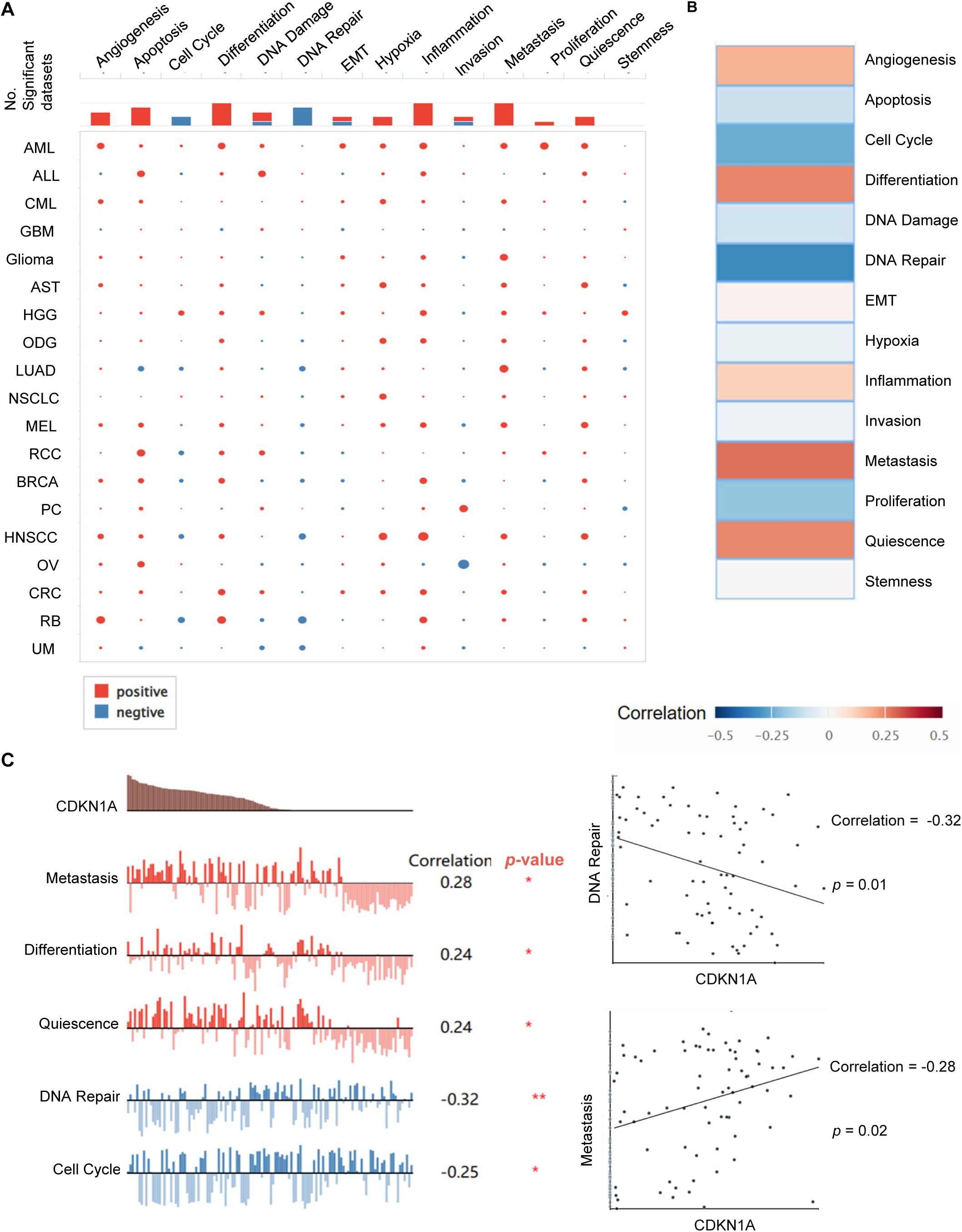
The functional relevance of CDKN1A across different cancers from CancerSEA. (A) Correlations between CDKN1A and biological activities in different cancers. (B) Functional relevance of CDKN1A in LUAD. Red plots suggesting positive correlations while blue plots indicating negative correlations. (C) CDKN1A is correlated with metastasis, differentiation and quiescence, DNA Repair and Cell Cycle in LUAD.

Next, we conducted cellular experiments to examine the biological functions of CDKN1A/p21 in cell proliferation. We performed transcriptomic RNA-sequencing upon sip21 transfection in H1299 to identify the downstream genes (Figure 10A). We observed 143 upregulated genes, 813 downregulated genes in total (Figure. 10B). GESA analysis of the RNA-seq reveals that p21 affects cell adhesion, apoptosis and EMT (Figure. 10C). We next validated the 10 potential target genes which are most up-or downregulated (Supplementary Figure 5A, 5B). Enforced p21 inhibits CCND3 expression and induces TNFRSF9 and NR6A1 expression at the mRNA level and protein level in H1299, A549, MCF-7 and BT474 cells, respectively (Supplementary Figure 5A, B, Figure. 10D). p21 overexpression dramatically reduced cell proliferation and clonogenic ability in lung cancer cells H1299, A549, in breast cancer cells MCF-7 and BT474, in prostate cancer cells Mia and PDC0034 (Figure 10E-10H). Conversely, knocking down of p21 promotes cell growth in lung cancer cells H1299 and breast cancer cells MCF-7 (Figure 10I, 10J). Furthermore, enforced p21 promotes cell apoptosis in lung cancer, breast cancer and prostate cancer (Figure 11). Additionally, p21 overexpression inhibits wound closure, whereas silenced p21 facilitates wound closure (Figure 12). Moreover, p21 overexpression facilitates the cell sensitivity to trametinib and cisplatin in H1299, A549, MCF-7 and BT474 cells, respectively (Supplementary Figure 5C, 5D). Cellular senescence, a hallmark of aging, refers to a state of stable and irreversible cell cycle arrest that occurs in response to various stressors, including DNA damage, oxidative stress, and oncogene activation [42]. Our study demonstrates that p21 induces cell senescence in lung cancer, breast cancer and prostate cancer cell lines (Figure 13).

**Figure 10.**
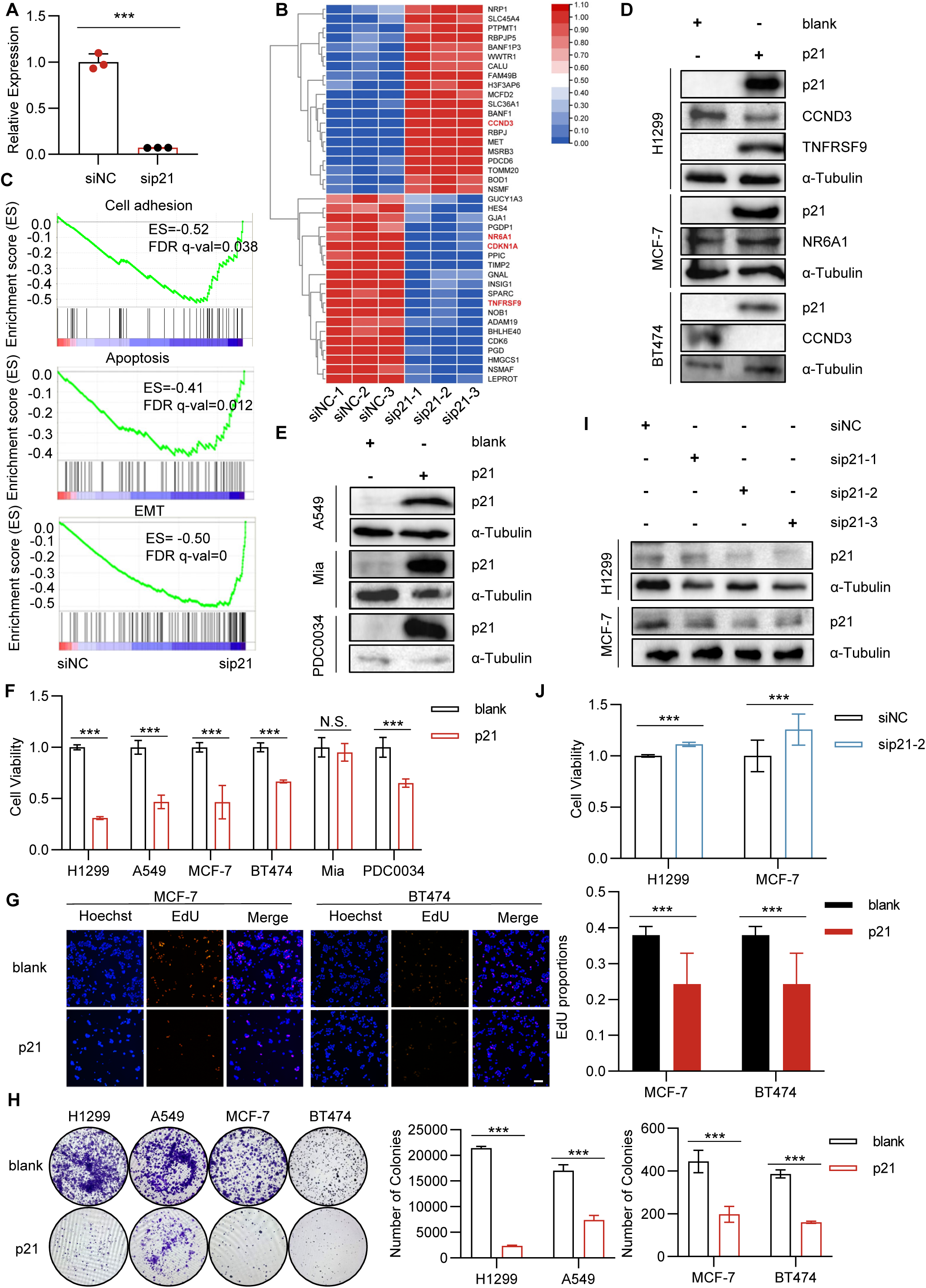
CDKN1A regulates cell proliferation. (A) qPCR result showing dramatic silencing of CDKN1A upon siRNA transfection in H1299 cells. (B) Heatmap showing the differentially expressed genes upon silenced p21 in H1299. (C) GESA analysis of the RNA-seq reveals that p21 affects cell adhesion, apoptosis and EMT. (D)Western blot revealing the dysregulated level of p21-modulated genes in different cell lines. (E-H) p21 overexpression (E) inhibits cell proliferation via CCK8 assay (F), EdU assay (G) and colony formation (H) in different cell lines. Scale bar, 100 μm. (I-J) Silenced p21 (I) inhibits cell growth in H1299 and MCF-7cells (J). *p < 0.05, **p < 0.01, ***p < 0.001.

**Figure 11.**
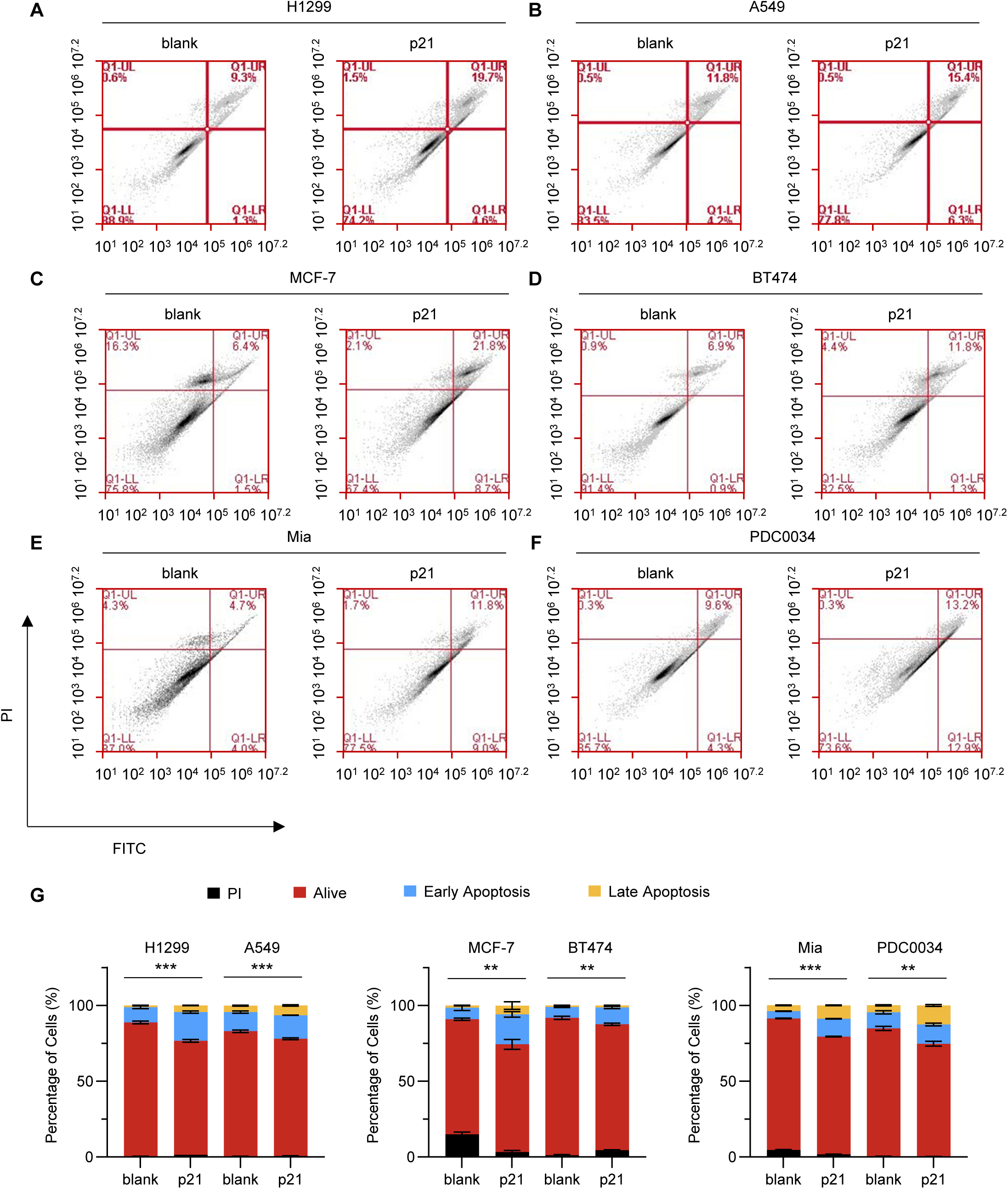
Representative Annexin-V plots indicating apoptosis distribution in multiple cells upon p21 overexpression in different cell lines.

**Figure 12.**
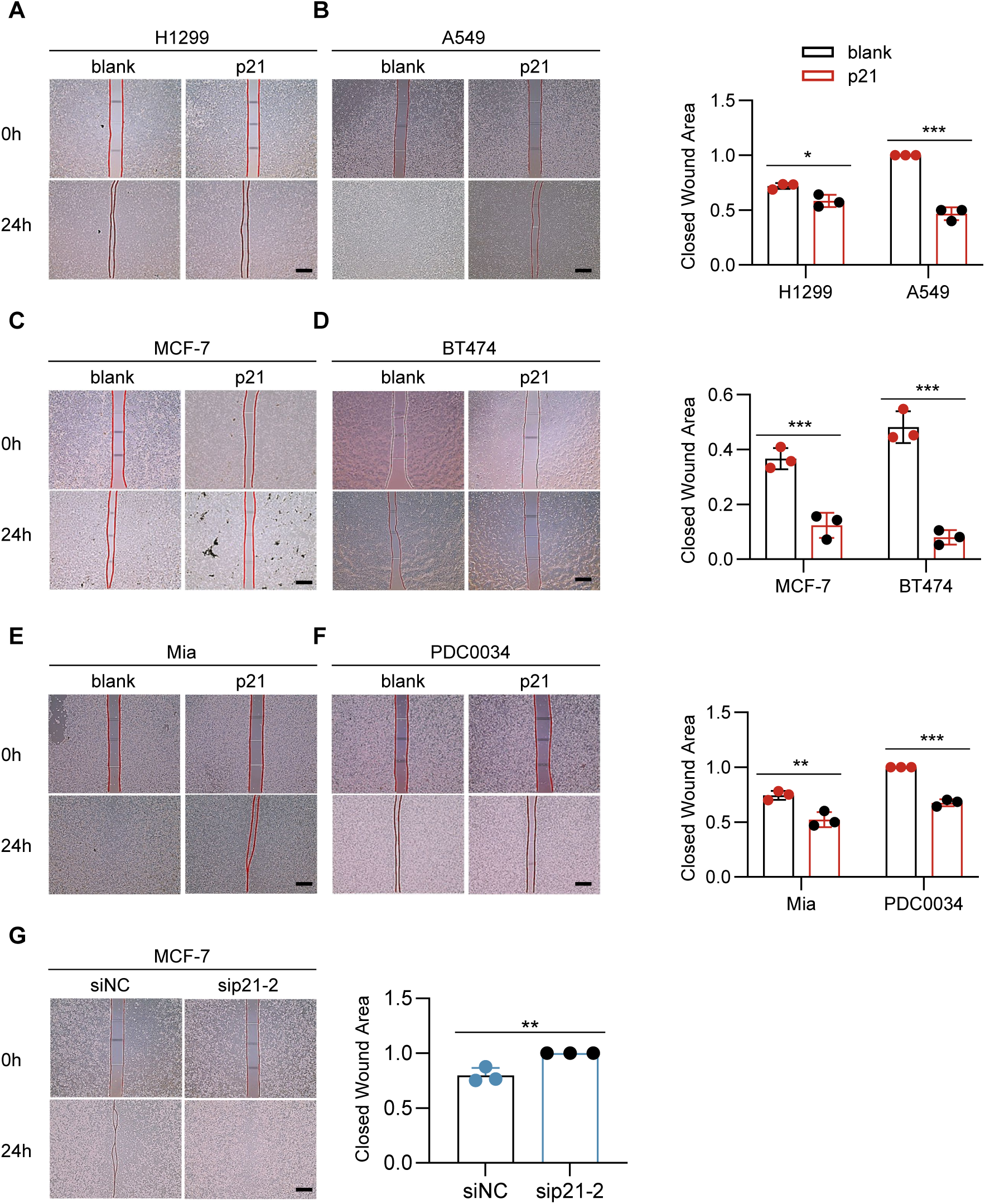
Scratch assay showing p21 modulates wound closure. Enforced p21 prevents wound closure (A-F), while silenced p21 enhances wound closure (G) in different cell lines. Scale bar, 100 μm. *p < 0.05, **p < 0.01, ***p < 0.001.

**Figure 13.**
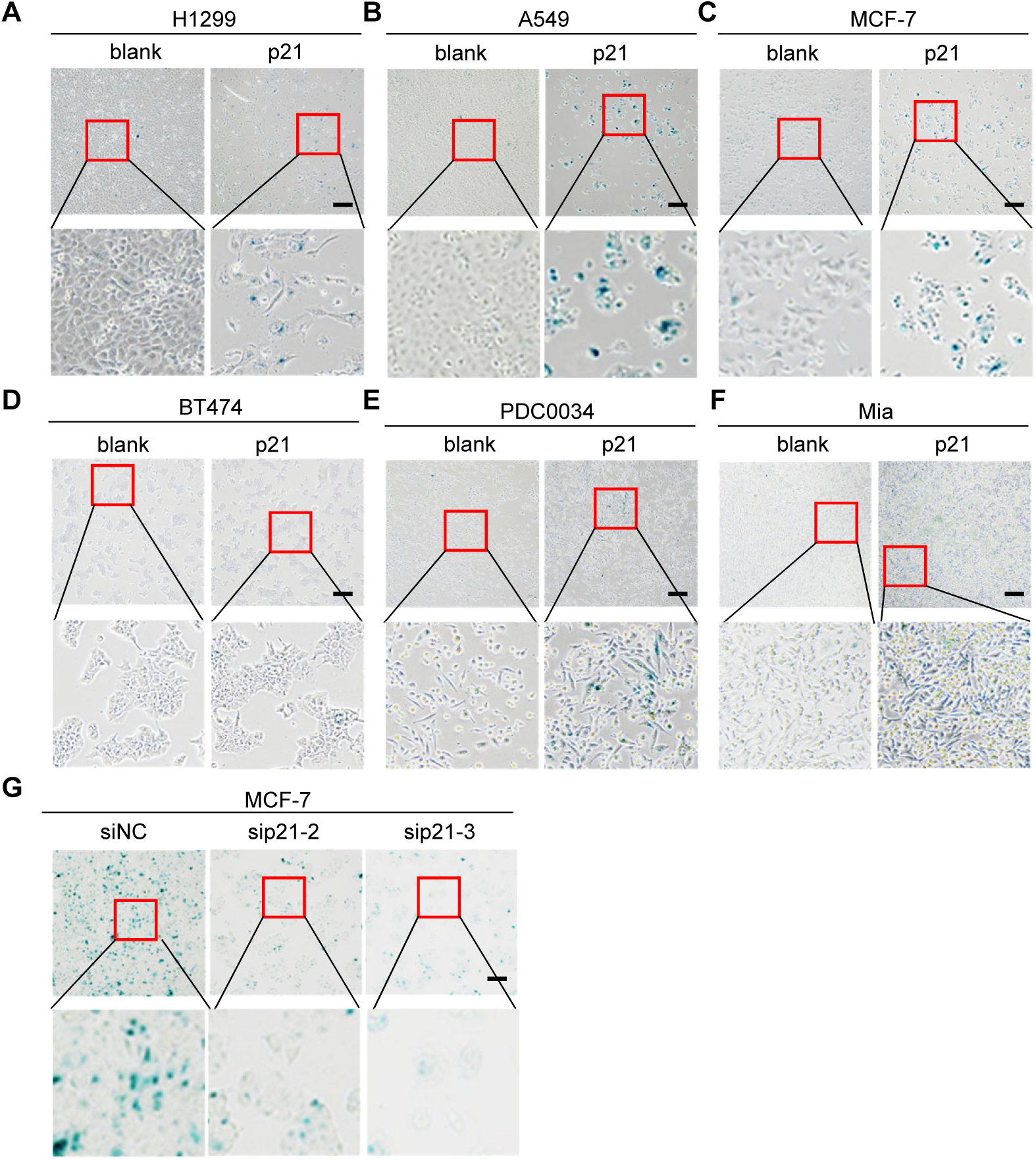
Senescence-associated-β-galactosidase (SA-β-gal) staining assay indicating overexpression of p21 promotes cell senescence (A-F), while inhibited p21 prevents cell senescence (G) in different cell lines. Scale bar, 100 μm.

In addition, we constructed a gene interaction network of CDKN1A using the GeneMANIA database. As shown in Supplementary Figure 6A, CDKN1A exhibits a strong physical interaction with PCNA, a key modulator in DNA replication and repair [41]. CDKN1A is also associated with cyclin-dependent kinases such as CDK6, CDK5, CDK4, CDK2, cyclin genes including CCNB1, CCNE2, CCND3, CCNA1 and others (Supplementary Figure. 6A). In addition, we observed that CDKN1A is closely associated with cancer driver genes and drug target genes in BRCA (Supplementary Figure. 6B), ccRCC (Clear cell renal cell carcinomas) (Supplementary Figure. 6C), EC (Endometrial cancer) (Supplementary Figure. 6D) and LUAD (Supplementary Figure. 6E). These findings suggest CDKN1A may affect tumorigenesis via these target genes.

### 3.8 Prognostic value of CDKN1A in cancer

In the end, we investigated the potential implication of CDKN1A as a prognostic marker in cancers. We evaluated its expression in patients across different tumor stage I, II, III and IV as well as in normal tissues via the UALCAN database. Compared to normal samples, CDKN1A level is lower in the late stages of BLCA (Supplementary Figure. 7A), BRCA (Supplementary Figure. 7B), COAD (Supplementary Figure. 7C), LUAD (Supplementary Figure. 7D), LUSC (Supplementary Figure. 7E), KICH (Supplementary Figure. 7F) and READ (Supplementary Figure. 7G), whereas the opposite trend is observed in CHOL (Supplementary Figure. 7H), HNSC (Supplementary Figure. 7I), KIRC (Supplementary Figure. 7J), KIRP (Supplementary Figure. 7K), THCA (Supplementary Figure. 7L) and CESC (Supplementary Figure. 7M), which is consistent with the CDKN1A mRNA levels in pan-cancer (Figure 2). However, we did not observe the significant differential expression in LIHC and ESCA (Supplementary Figure. 7N, 7O). Furthermore, univariate analysis shows that LUSC tumorigenesis is associated with CDKN1A at the different stages (Table IV). ROC curves were used to assess the relevance of genetic markers on diagnostic accuracy [43]. Different area under the curve (AUC) thresholds have been identified to represent high diagnostic accuracy (AUC: 1.0-0.9), moderate diagnostic accuracy (AUC: 0.9-0.7), or low diagnostic accuracy (AUC: 0.7-0.5). CDKN1A has relatively moderate sensitivity and specificity for PAAD (Supplementary Figure. 8A), READ (Supplementary Figure. 8B) and LUSC (Supplementary Figure. 8C), suggesting that CDKN1A may serve as a prognostic biomarker. However, the low diagnostic accuracy of CDKN1A for BRCA (Supplementary Figure. 8D), BLCA (Supplementary Figure. 8E), COAD (Supplementary Figure. 8F), LUAD (Supplementary Figure. 8G) and UVM (Supplementary Figure. 8H) suggests that it is unsuitable as a prognostic marker for these cancers.

We next investigated the relationship between CDKN1A expression and the prognostic values in pan-cancer, including OS, DSS, DFI and RFS. Using the PrognoScan database, we found that CDKN1A expression is substantially associated with OS in 8 cancer types (Supplementary Figure. 9). In particular, high expression of CDKN1A is associated with better clinical survival outcomes in bladder cancer (Supplementary Figure. 9A), blood cancer (Supplementary Figure. 9B), colorectal cancer (Supplementary Figure. 9C), breast cancer (Supplementary Figure. 9D-F), eye cancer (Supplementary Figure. 9G) and ovarian cancer (Supplementary Figure. 9H-I), whereas enforced CDKN1A is correlated with worse outcomes in lung cancer (Supplementary Figure. 9J-K) and brain cancer (Supplementary Figure. 9L).

Based on the Kaplan-Meier plotter database, elevated CDKN1A is associated with poor OS in BLCA (Supplementary Figure. 10A, Left), BRCA (Supplementary Figure. 10B, Left), ESCA (Supplementary Figure. 10C, Left), HNSC (Supplementary Figure. 10D, Left) and LUSC (Supplementary Figure. 10E, Left), but CDKN1A correlates with protective survival in KIRP (Supplementary Figure. 10F, Left), LIHC (Supplementary Figure. 10G, Left) and KIRC (Supplementary Figure. 10H, Left). RFS results show a consistent trend of clinical outcome, but we observed beneficial outcomes in BRCA (Supplementary Figure. 10B, Right) and LUSC (Supplementary Figure. 10E, Right).

CDKN1A is a protective factor in KIRC and KIRP, and a risk factor in DLBC, GBM, LGG, LUSC, MESO, PAAD and UVM (Figure. 14A). In addition, CDKN1A expression is substantially associated with DSS in seven cancers and PFI in five cancers. In the DSS analysis, CDKN1A is a protective factor in KIRC, KIRP and UCEC, whereas a risk factor in LUSC, PAAD, THYM and UVM (Figure. 14B), while the PFI analysis shows that CDKN1A is a risk factor in GBM, LUSC, and a protective factor in KIRC, KIRP and UCEC (Figure. 14C). In the RFS study, cox regression analysis of 33 tumors show that CDKN1A expression is significantly associated with RFS and acts as a protective factor in KIRP (Figure. 14D). The OS survival analysis of CDKN1A shows that patients with high CDKN1A gene expression have better survival in KIRC (Figure. 14E), whereas high-expressed CDKN1A is associated with worse outcomes in LGG and PAAD, receptively (Figure. 14F, G). In summary, these data suggest the prognostic role of CDKN1A in cancer.

**Figure 14.**
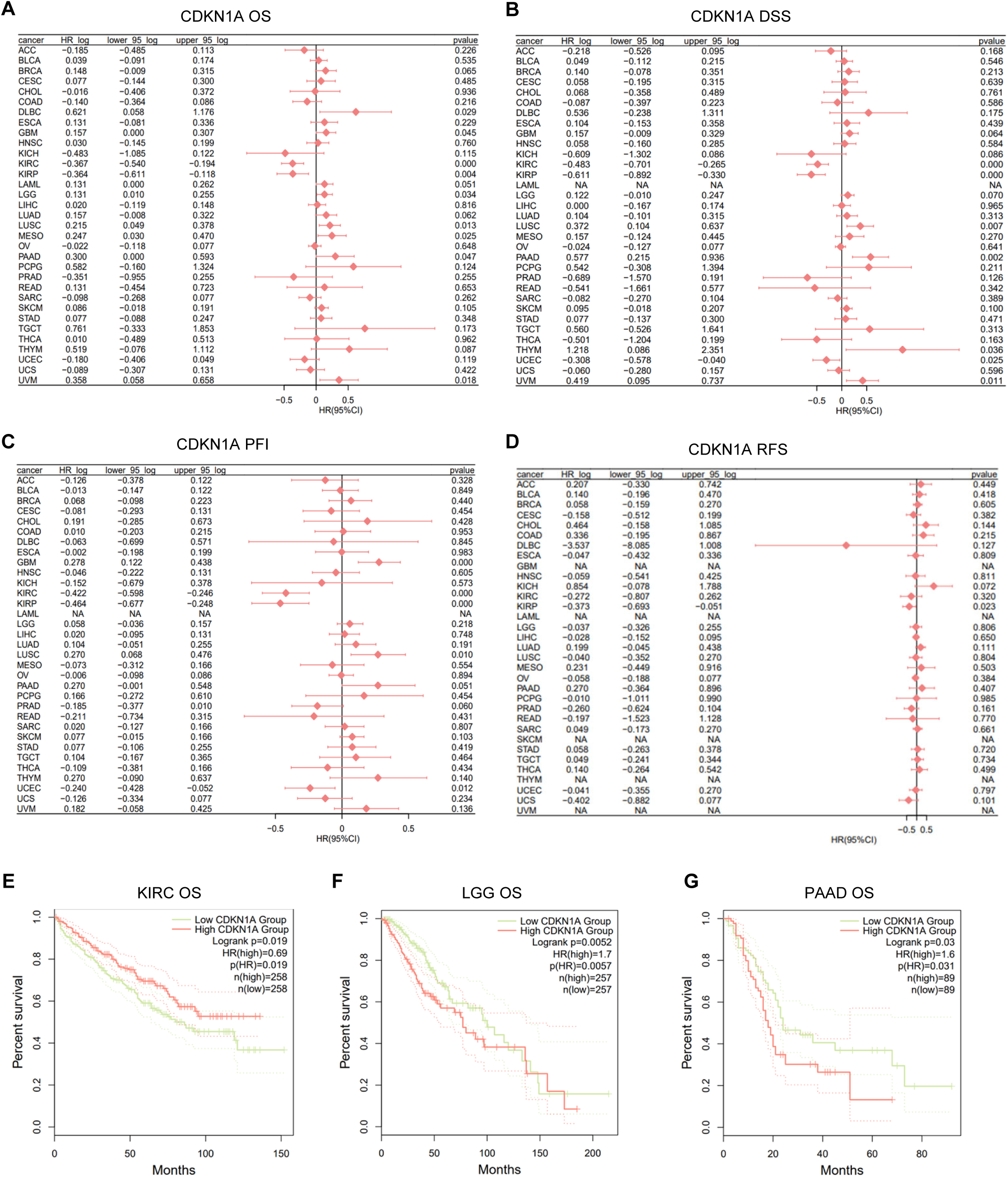
CDKN1A is associated with patient outcomes. (A-D) The forest plot showing the relationship of CDKN1A expression with overall survival (A), disease-specific survival (B), progression-free interval (C) and disease-free survival (D). (E-G) Kaplan-Meier analyses indicating the association between CDKN1A expression and OS in KIRC (E), LGG (F) and PAAD (G).

## 4. Discussion

Cancer cells exhibit genetic instability, undergoing multiple genetic and epigenetic changes as they continuously evolve in response to selective pressures. Cell proliferation, motility, apoptosis, senescence, motility and survival are regulated by multiple genomic alternations [3]. Cell cycle is a hallmark of human cancer since tumor cells often harbor DNA damage that directly regulate their cell cycles [44]. Recent efforts have identified that cyclins and the catalytic partners, termed as cyclin-dependent kinase (CDKs), are key players in the transition between different phases of the cell cycle [45, 46]. Specifically, CDK1/2/4/6 facilitate cell cycle progression by phosphorylating retinoblastoma protein (RB) and silencing of TP53, thereby removing the G1 checkpoint restriction and promoting entrance into S phase during cancer development [47]. CDKs are negatively regulated by two distinct families of cyclin-dependent kinase inhibitors (CKIs): the inhibitor of kinase (INK) family, comprising four structurally related proteins (p16^INK4A^, p15^INK4B^, p18^INK4C^ and p19^INK4D^), and the CDK-interacting protein/kinase inhibitory protein (CIP/KIP) family, which comprises three proteins (p21^Cip1^, p27^Kip1^ and p57^Kip2^) [45]. p21, encoded by gene CDKN1A, functions as a bone-fide cell cycle silencer and anti-proliferative effector by triggering cell cycle arrest in G1 and G2 phases of tumor cells. Previous studies have identified that p21 as a transcriptional target of p53, participating in a wide range of events through both p53-dependent and p53-independent pathways [11, 48]. Dong et al., recently reported that Lamin B2 (LMNB2) promotes cell proliferation and metastasis via affecting p21-mediated cell cycle progression in colorectal cancer [49]. Additionally, lncRNA such as PCAT-1 and lncRNA MSTO2P can also enhance drug resistance and proliferation by repressing p21 in gastric cancer and colorectal cancer, respectively [50, 51].

In the present study, we systemically analyzed the copy number variant, expression, prognostic values and biological functions of CDKN1A/p21 in diverse cancer types. We did not observe a correlation between CNV and expression of CDKN1A. At the mRNA level, CDKN1A is lowly expressed in BLCA, BRCA, COAD, KICH, LUAD, LUSC, PRAD, READ and STAD, in contrast, highly expressed in CHOL, HNSC, KIRC, KIRP and THCA compared to corresponding normal tissues. At the protein level, CDKN1A/p21 is lower in colon cancer, breast cancer and prostate cancer. Interestingly, we observed an increased CDKN1A/p21 protein level in lung cancer, implying an alternative mechanism occurs in p21 post-transcription regulation. In combination with high CDKN1A expression, it presents a favorite and/or adverse influence on clinical outcomes in pan-cancer, suggesting CDKN1A may characterize a dual role in the progression of cancers. Emerging studies have identified that CDKN1A/p21 as a dual regulator depending on cell types, cancer types, cellular localizations, the presence or absence of p53, and genotoxic stress. A recent breakthrough revealed that enforced p21 inhibits CRL4-CDT2 ubiquitin ligase, promotes deregulated origin licensing and replication stress, eventually stimulating genomic instability, aggressiveness and chemoresistance [52]. *In vitro* and *in vivo* studies gained contradictory data for CDKN1A/p21, presenting either chemosensitization or chemoresistance in different cancer types. For example, CDKN1A/p21 deletion enhances cell sensitivity to cisplatin in human colon cancer HCT-116 cells and murine embryonic fibroblasts (MEFs) [53]. We found that ectopic p21 enhances the cell sensitivity to trametinib and cisplatin in lung cancer and breast cancer. Given the dual roles of p21 in certain cellular contexts, targeting p21 for therapeutic intervention is a promising but challenging task. We then examined the biological functions of CDKN1A in diverse cancers. The gene signature reveals that CDKN1A is associated different crucial signaling pathways in cancer, including KRAS signalling and MYC signalling in BLCA, COAD, LUAD and LUSC. KRAS, a principal isoform of RAS, is associated with over one-third of all cancers by stimulating multiple effectors including RAF and PI3K [54]. Wang et al., identified that co-administration of inhibitors of polo-like kinase 1 and RhoA/Rho kinase (ROCK) increases the expression of p21, therefore rendering KRAS G12D induced lung cancer [55]. Morton reported that LKB1 haploinsufficiency can cooperate with oncogenic KRAS G12D to lead tumorigenesis by repressing CDKN1A/p21-dependent cell cycle arrest in PDAC [56]. MYC is an oncogenic transcription factor that directly binds to the E-box in the genes which are involved in the cell cycle and key processes of cell growth, apoptosis, differentiation and metabolism [57]. We observed that MYC overexpression downregulates p21 in different cancer. MYC is a well-established transcriptional repressor of p21. Jung and Hermeking discovered that MYC dives a transcriptional cascade by inducing the transcription factor AP4 (TFAP4), which binds to recognition motifs near the p21 promoter and mediates its repression [58, 59]. Seoane reported that MYC recruits to the p21 promoter, displacing p53 and inhibiting p21 transcription [60]. In addition, mice lacking PUMA, NOXA and p21, or bearing mutations in p53, accelerate MYC-driven lymphoma development *in vivo* [61]. In summary, our findings suggest that CDKN1A may participate in a series of signalling pathways therefore affecting tumorigenesis.

The tumor microenvironment influences tumor growth, metastatic and therapy response. TME-mediated immunosuppression often impairs beneficial responses, complicating the design of new therapies [62]. To further validate the potential relationship between CDKN1A and immune microenvironment, we systematically analyzed the correlation between CDKN1A and tumor immunity. Our results showed that CDKN1A is closely associated with infiltration of various immune cells, including B cells, CD8+ T cells, CD4+ T cells, macrophages, neutrophils and dendritic cells, suggesting that CDKN1A may be involved in the immune regulation in multiple tumors, which may ultimately affect patient survival. Gene signature analysis indicates the enrichment of immunological pathways such as IL6/JAK/STAT3 signalling, IL17 signalling and TNF signalling in diverse tumors. Thoma found that p21 knockout mice accumulate abnormal amounts of CD4+ memory cells and develop a loss of tolerance towards nuclear antigens [63]. Lawson and authors reported that ablation of CDKN1A/p21 leads to active Fas/FasL-induced T cell death, enhancing survival and reduces kidney disease in p21−/− BXSB mice [64]. However, the function of CDKN1A/p21 in cancer immunity remains to be fully elucidated. The heterogeneity among individual cells have profound influences in both unicellular and multicellular organisms.

Recently advanced scRNA-seq enable unbiased, high-throughput and high-resolution transcriptomic analysis of individual cells [65]. Our scRNA-seq analysis revealed significant CDKN1A distribution in cDC3, AT2, and macrophage cells, highlighting CDKN1A as a potential factor in tumor immunological therapy.

We conducted functional experiments and found that p21 overexpression leads to a significant reduction in proliferative capacity via CCK8, colony formation, EdU and wound scratch assay, and promotes cell apoptosis and cell senescence in different cancer types. Meanwhile, silenced p21 facilitates cell growth and wound closure, prevent cell senescence in different cancer cell lines.

In summary, this study provides a preliminary basis for understanding the function of CDKN1A in pan-cancer, but its specific biological role and mechanism require further experimental validation. In addition, the current study may have some shortcomings with inconsistent results, possibly due to the data obtained from multiple bioinformatics databases.

## 5. Conclusion

Our study explored the expression levels of CDKN1A in different cancer types, suggesting that CDKN1A could serve as a potential prognostic marker. Our findings reveal that CDKN1A exerts a significant inhibitory effect on cancer by suppressing cell proliferation and migration, inducing cell senescence, and promoting apoptosis, which lays the foundation for further investigation of the prognostic and therapeutic potential of CDKN1A.

## 7. Author Contributions

WYZ, QLM, WRL, HHZ, LHZ, YNX, YYR, KXY, YHL and LS conceptualized the review, performed the literature search and wrote the manuscript. YHL and LS revised each step of the work and are responsible for the final revision. All authors read and approved the final manuscript.

## 8. Data Availability Statement

The datasets used to support the results of this study are available from the corresponding author upon request.

## 10. ORCID

Lei Shi https://orcid.org/0000-0003-4027-2396.

---

**Supplementary Figure 1.**
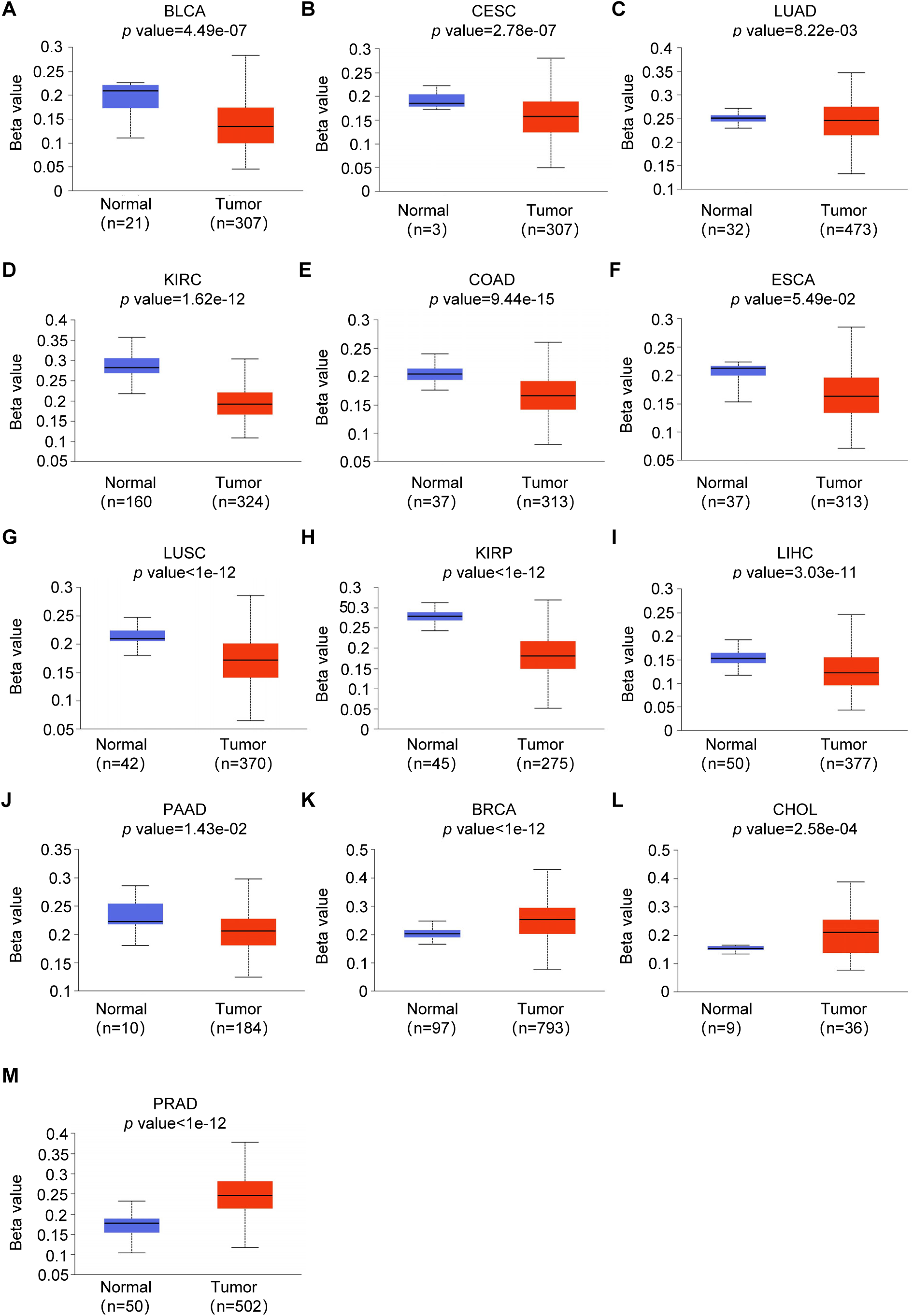
Histogram bars showing the hyper-or hypomethylation of the CDKN1A promoter region in tumors compared to normal tissues in BLCA (A), CESC (B), LUAD (C), KIRC (D), COAD (E), ESCA (F), LUSC (G), KIRP (H), LIHC (I), PAAD (J), BRCA (K), CHOL (L), PRAD (M).

**Supplementary Figure 2.**
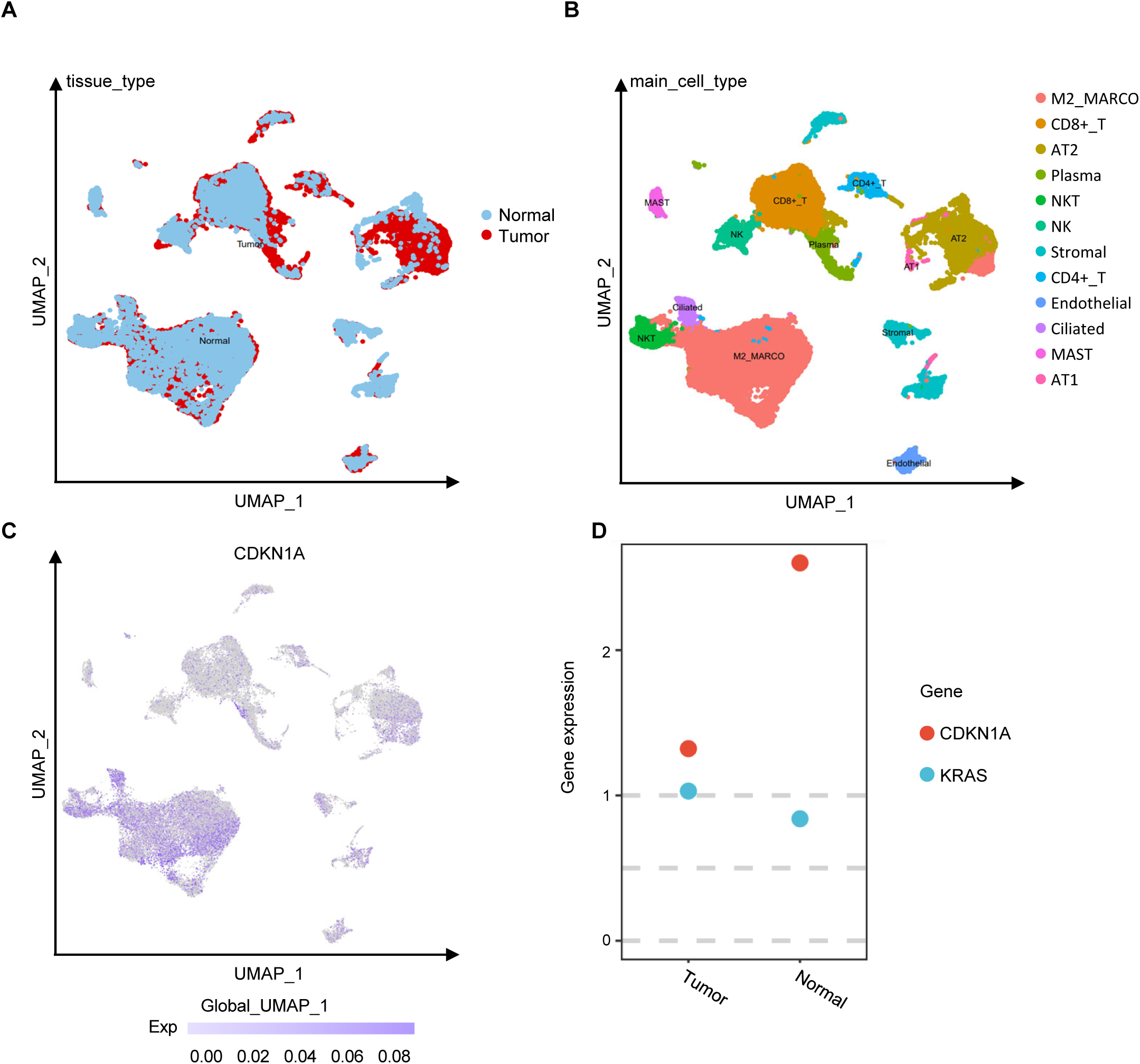
scRNA analysis of CDKN1A distribution in LUAD. (A-C) scRNA-seq revealing the CDKN1A gene distribution in normal/tumor samples (A), cell subpopulations (B) and CDKN1A gene expression in LUAD (C). (D) The expression of CDKN1A and KRAS in normal/tumor cohorts.

**Supplementary Figure 3.**
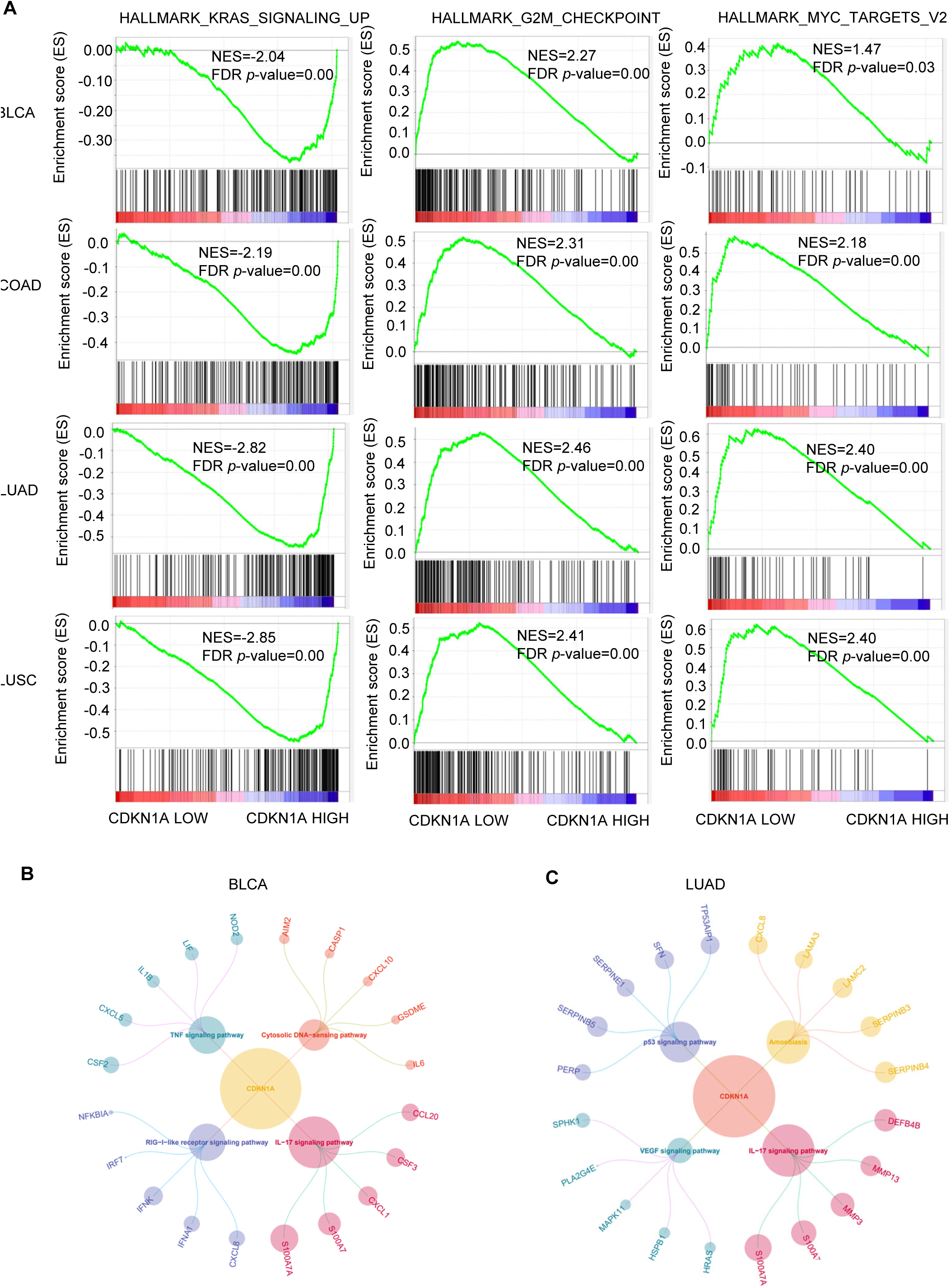
Gene signature analysis in different cancers. (A) “KRAS”, “G2M-CHECKPOINT” and “MYC” signalling pathways are enriched in LUAD, LUSC, BLCA and COAD. (B, C) Gene Signature analysis in BLCA (B) and LUAD (C).

**Supplementary Figure 4.**
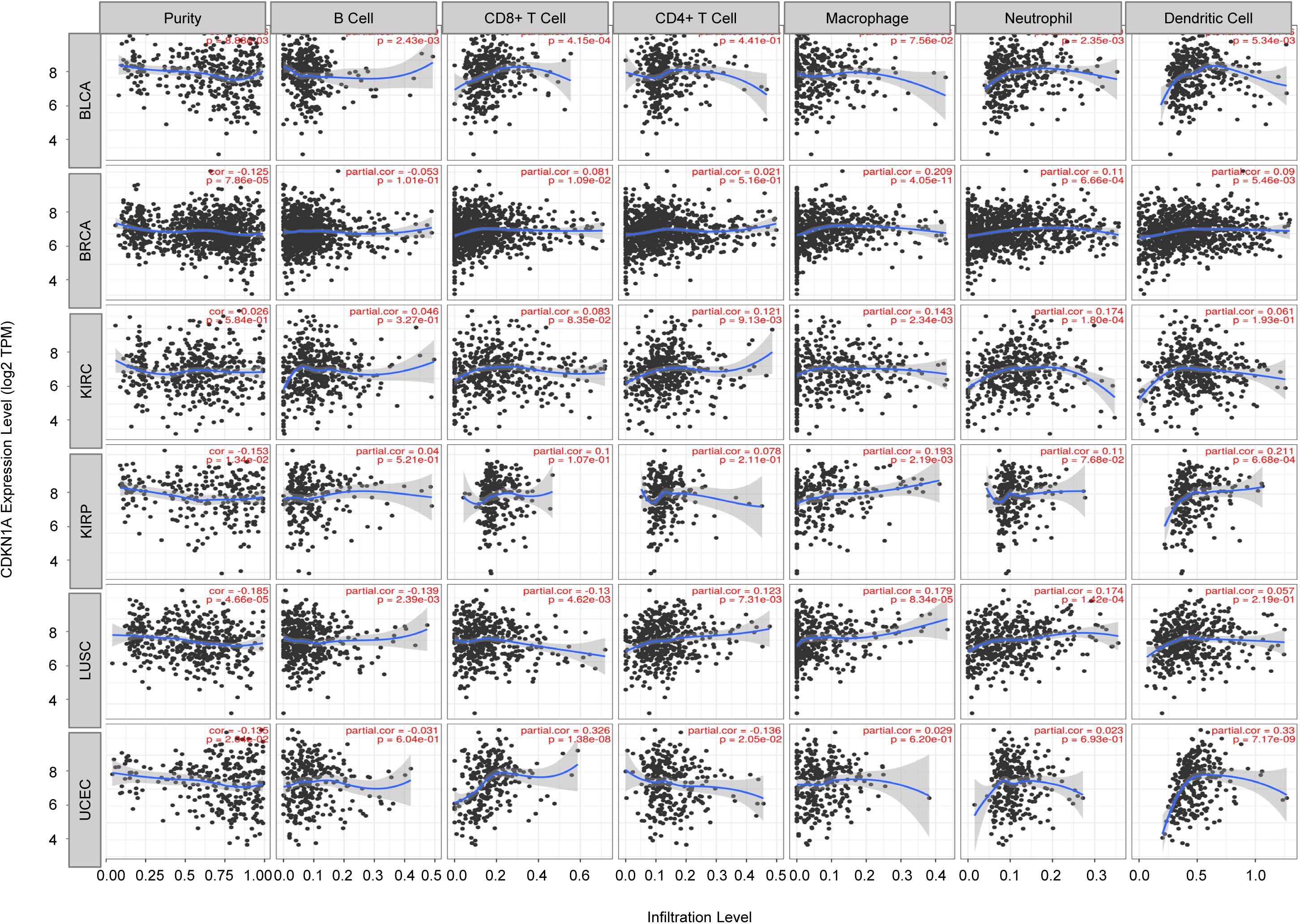
Correlation between CDKN1A levels and immune cell infiltration in different cancer types via TIMER database.

**Supplementary Figure 5.**
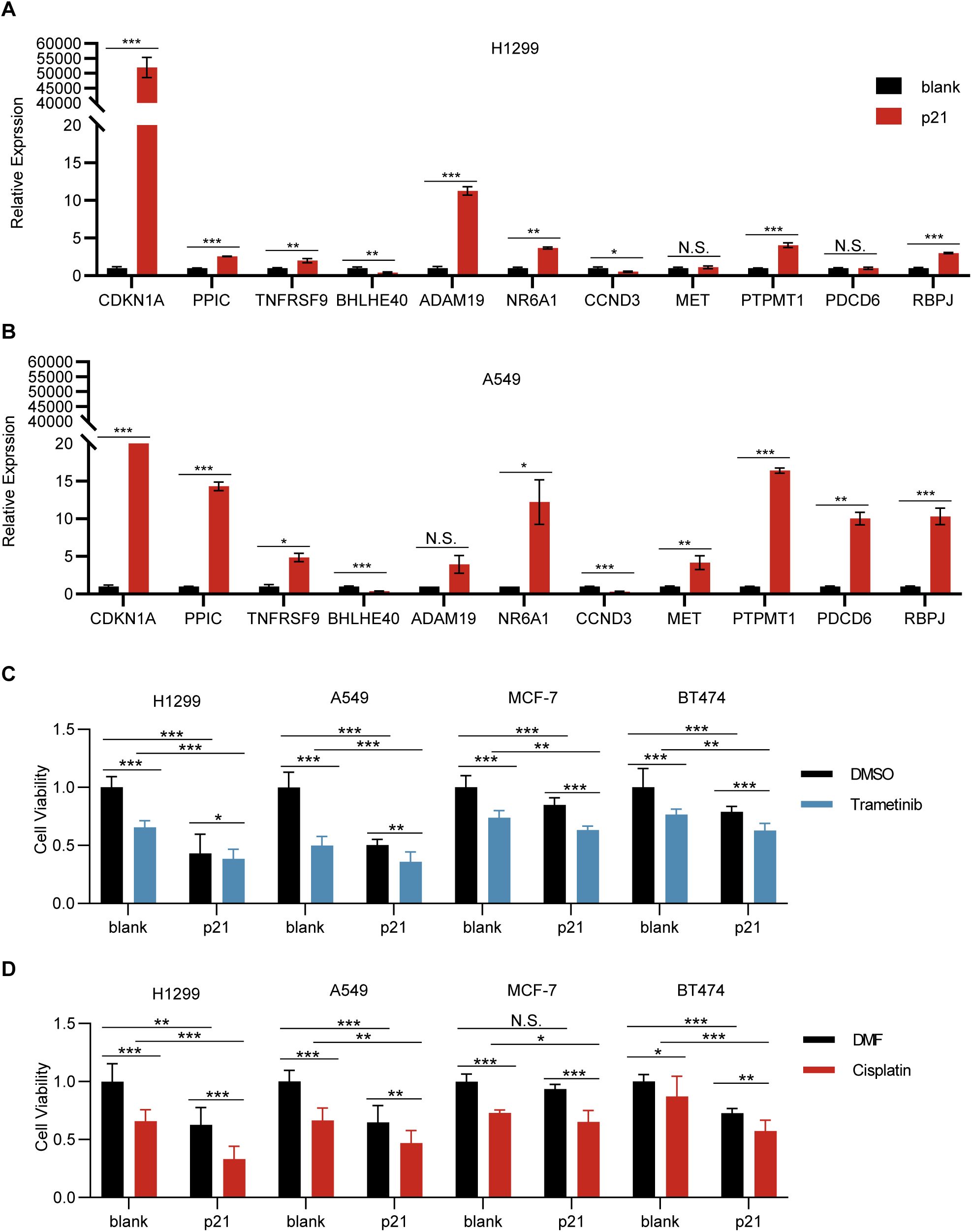
p21 regulates gene expression and promotes drug sensitivity. (A, B) qPCR showing dysregulated genes after p21 overexpression in H1299 (A) and A549 (B). (C-D) p21 OE sensitize cells to Trametinib (C) and Cisplatin (D) treatment in H1299, A549, MCF-7 and BT474 cells, respectively. Trametinib, 5Μm; Cisplatin 5μM. ∗ *p* < 0.05, ∗∗ *p* < 0.01, and ∗∗∗ *p* < 0.001.

**Supplementary Figure 6.**
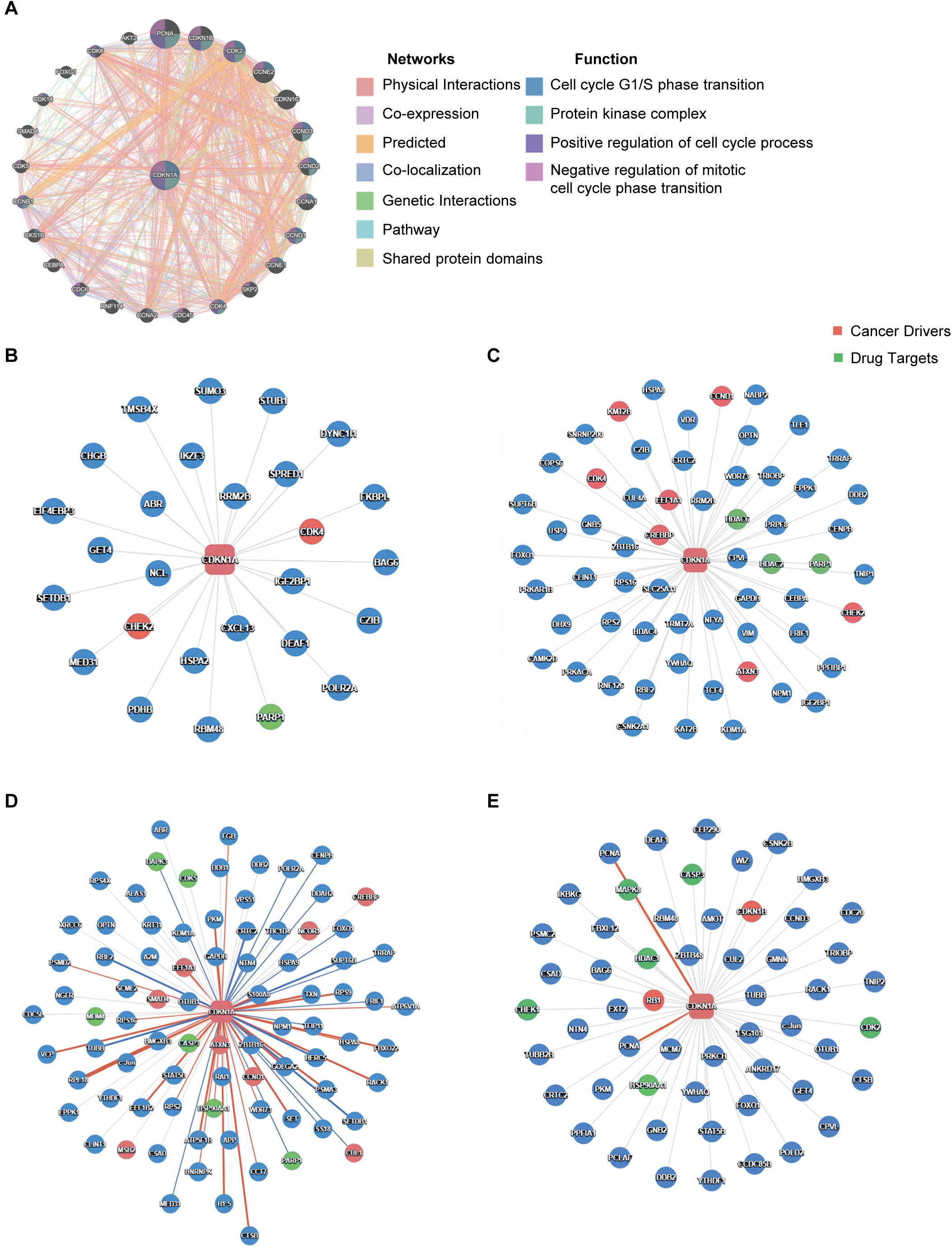
Gene network analysis of CDKN1A in pan-cancer. (A) The gene–gene interaction network of CDKN1A from GeneMANIA. (B-E) The p21-interacting protein network In BRCA (B), ccRCC (C), EC (D) and LUAD (E). ∗ *p* < 0.05, ∗∗ *p* < 0.01, and ∗∗∗ *p* < 0.001.

**Supplementary Figure 7.**
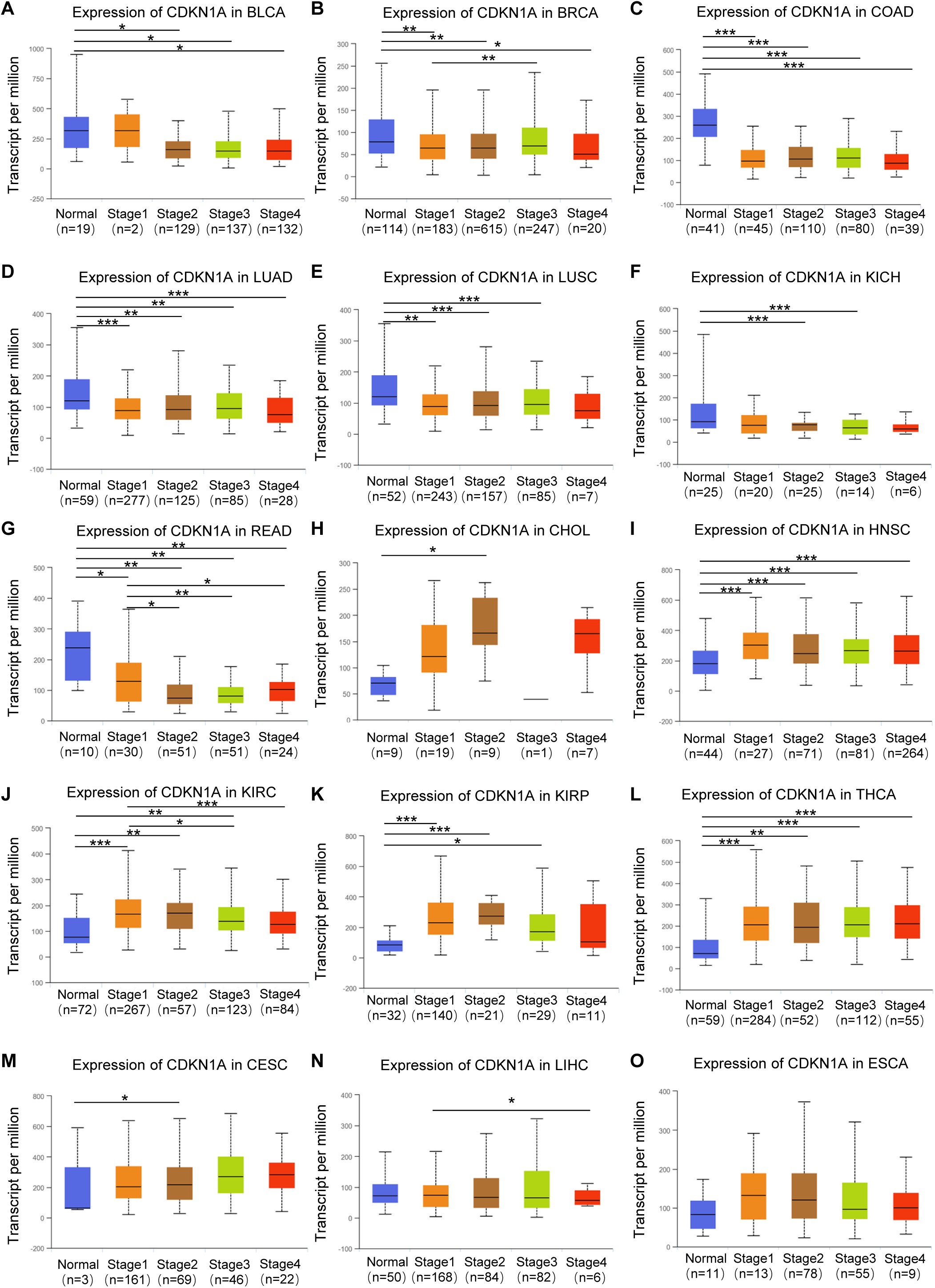
CDKN1A expression in different pathological stages of cancers. The expression level of CDKN1A in different tumor stage of BLCA (A), BRCA (B), COAD (C), LUAD (D), LUSC (E), KICH (F), READ (G), CHOL (H), HNSC (I), KIRC (J), KIRP (K), THCA (L), CESC (M), LIHC (N) and ESCA (O). ∗ *p* < 0.05, ∗∗ *p* < 0.01, and ∗∗∗ *p* < 0.001.

**Supplementary Figure 8.**
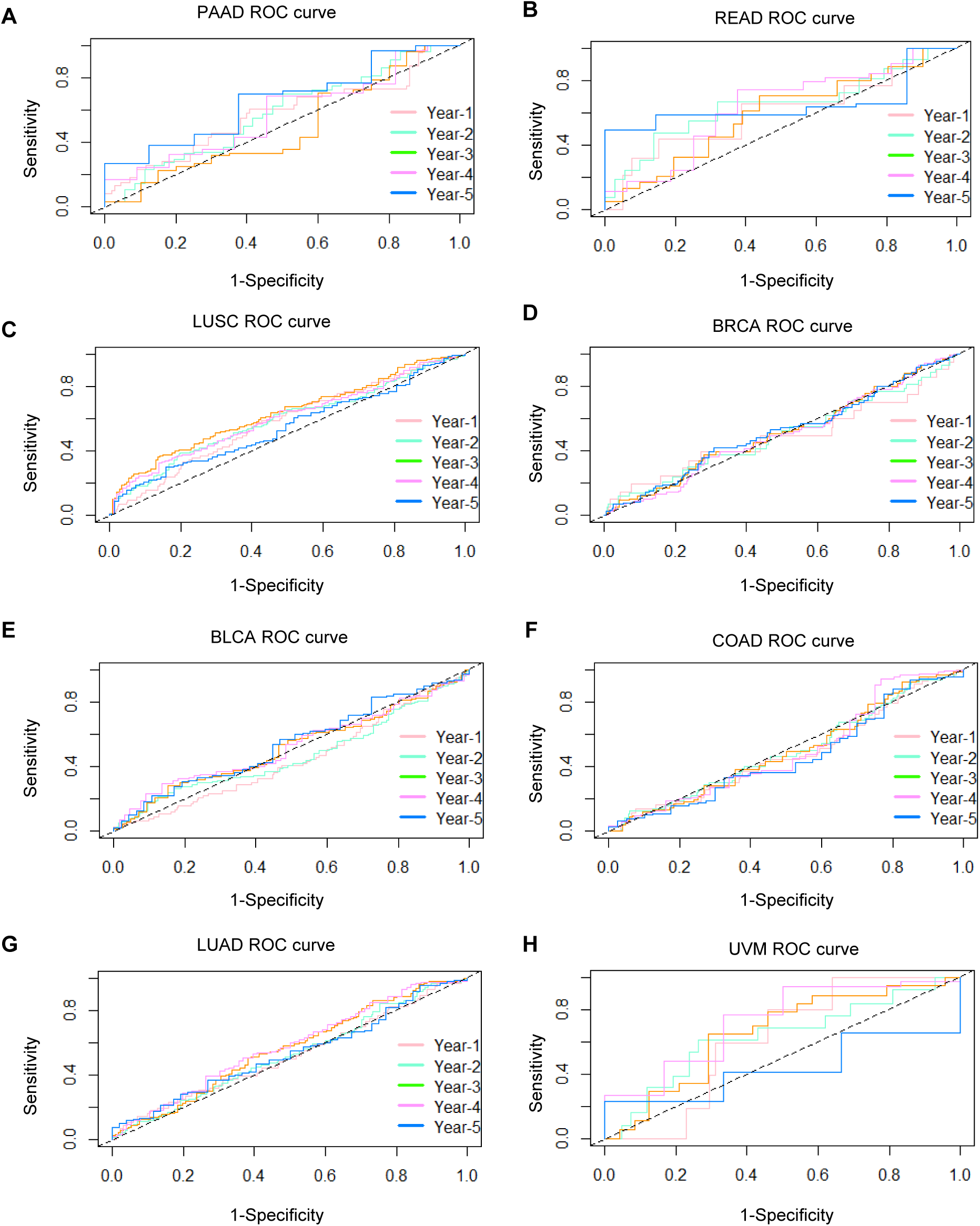
ROC curve displaying the relatively sensitivity and specificity of CDKN1A with cancer. ROC curves showing the predictive efficacy of CDKN1A with PAAD (A), READ (B), LUSC (C), BRCA (D), BLCA (E), COAD (F), LUAD (G), UVM (H) based on the TCGA cohort.

**Supplementary Figure 9.**
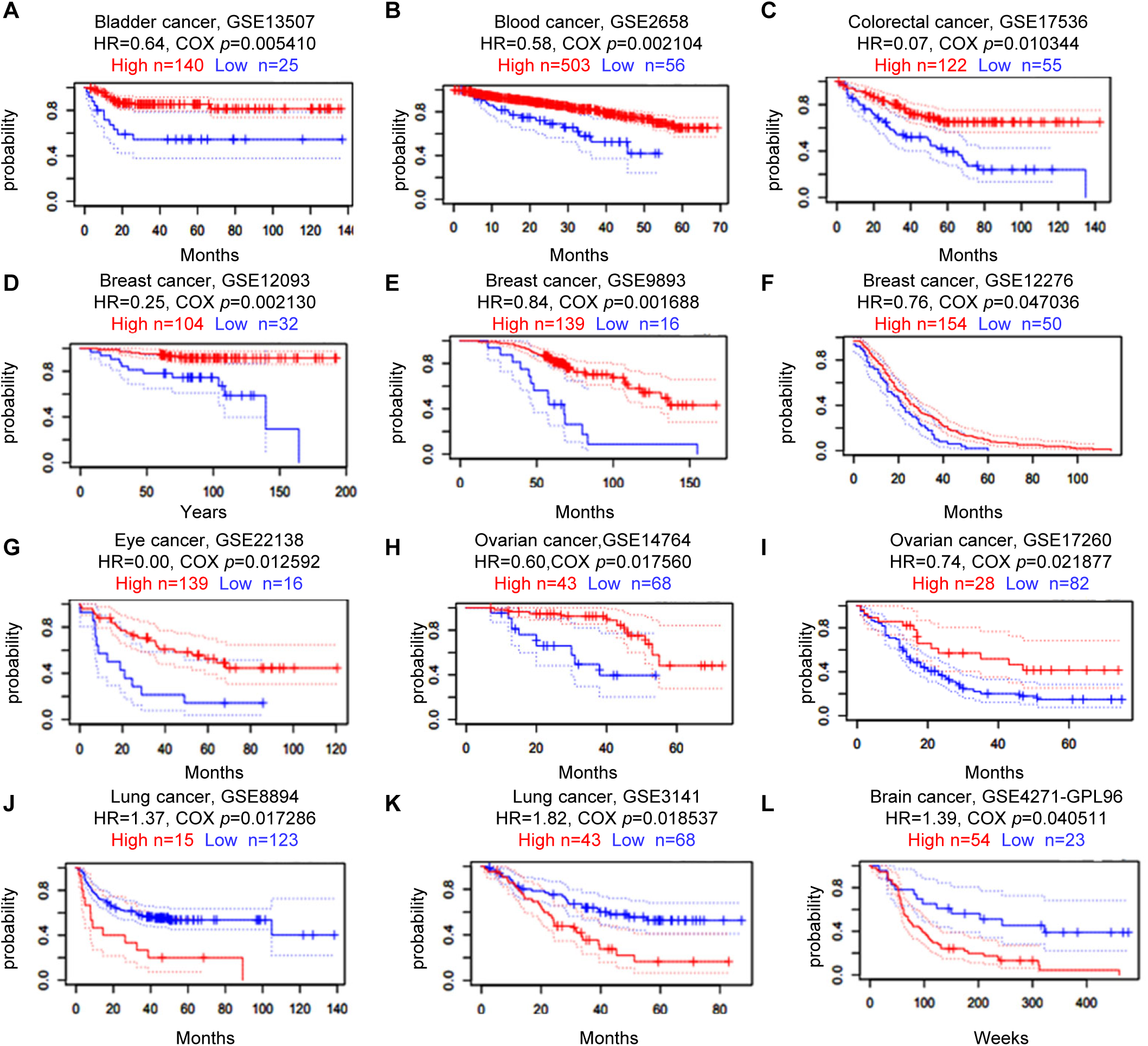
PrognoScan database showing the prognostic values of CDKN1A in 8 cancer types. High CDKN1A is associated with beneficial survival in bladder cancer (A), blood cancer (B), colorectal cancer (C), breast cancer (D-F), eye cancer (G) and ovarian cancer (H, I), whereas with adverse clinical outcomes in lung cancer (J, K) and brain cancer (L).

**Supplementary Figure 10.**
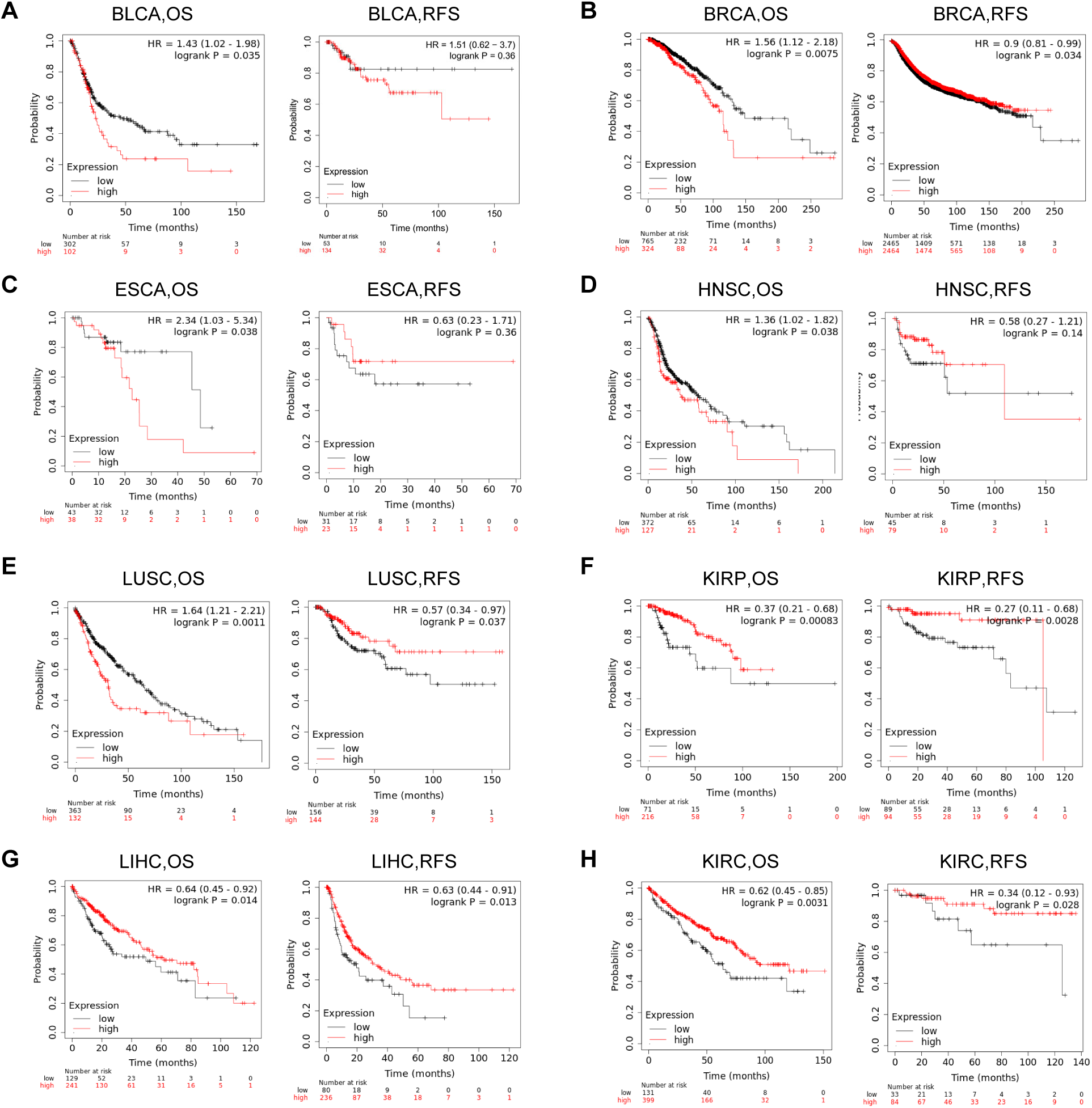
Kaplan–Meier survival curves indicating the clinical outcomes between high and low expression of CDKN1A in different cancer types. The OS and RFS survival analysis of CDKN1A in BLCA (A), BRCA (B), ESCA (C), HNSC (D), LUSC (E), KIRP (F), LIHC (G), KIRC (H).

**TABLE I.**
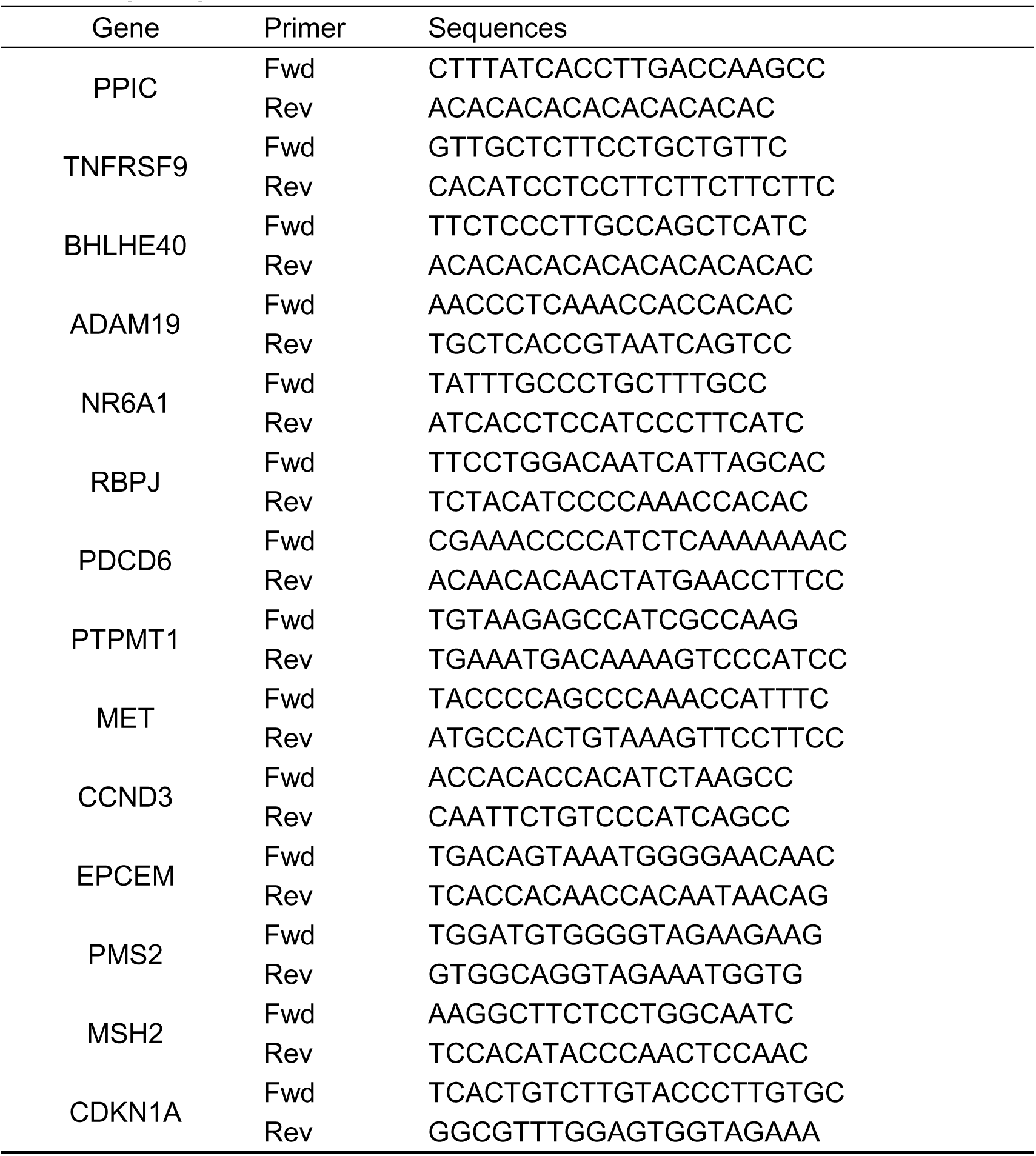
qPCR primers.

**TABLE II.** CDKN1A expression in different cancer type via UALCAN database.

| Cancer Type | Normal |  | Tumor |  | *p* |
| --- | --- | --- | --- | --- | --- |
|  | number | Median | number | Median |  |
| BLCA | 19 | 8.332 | 408 | 7.32 | 1.63E-02 |
| BRCA | 114 | 6.435 | 1097 | 6.142 | 5.46E-03 |
| CESC | 3 | 6.076 | 305 | 7.909 | 7.87E-01 |
| CHOL | 9 | 6.063 | 36 | 7.22 | 5.50E-02 |
| COAD | 41 | 8.055 | 286 | 2.798 | 1.00E-12 |
| ESCA | 11 | 6.461 | 184 | 6.844 | 6.26E-01 |
| GBM | 5 | 4.931 | 156 | 6.946 | 1.21E-01 |
| HNSC | 44 | 7.526 | 520 | 8.079 | 1.55E-05 |
| KICH | 25 | 7.06 | 67 | 6.317 | 3.80E-02 |
| KIRC | 72 | 6.338 | 533 | 7.343 | 7.63E-05 |
| KIRP | 32 | 6.431 | 290 | 8.024 | 4.82E-06 |
| LIHC | 50 | 6.234 | 371 | 6.326 | 9.78E-01 |
| LUAD | 59 | 7.003 | 515 | 6.561 | 1.85E-04 |
| LUSC | 52 | 7.446 | 503 | 6.964 | 7.56E-04 |
| PAAD | 4 | 7.283 | 178 | 7.139 | 2.85E-01 |
| PRAD | 52 | 6.322 | 497 | 5.938 | 8.64E-03 |
| PCPG | 3 | 6.449 | 179 | 7.133 | 4.77E-01 |
| READ | 10 | 8.011 | 166 | 6.527 | 1.12E-02 |
| SARC | 2 | 5.923 | 260 | 7.133 | 2.53E-01 |
| THCA | 59 | 6.308 | 505 | 7.797 | 1.96E-07 |
| THYM | 2 | 5.901 | 120 | 4.682 | 3.43E-01 |
| STAD | 34 | 7.317 | 415 | 6.492 | 2.30E-03 |
| UCEC | 35 | 6.64 | 546 | 6.512 | 6.46E-01 |

**TABLE III.**
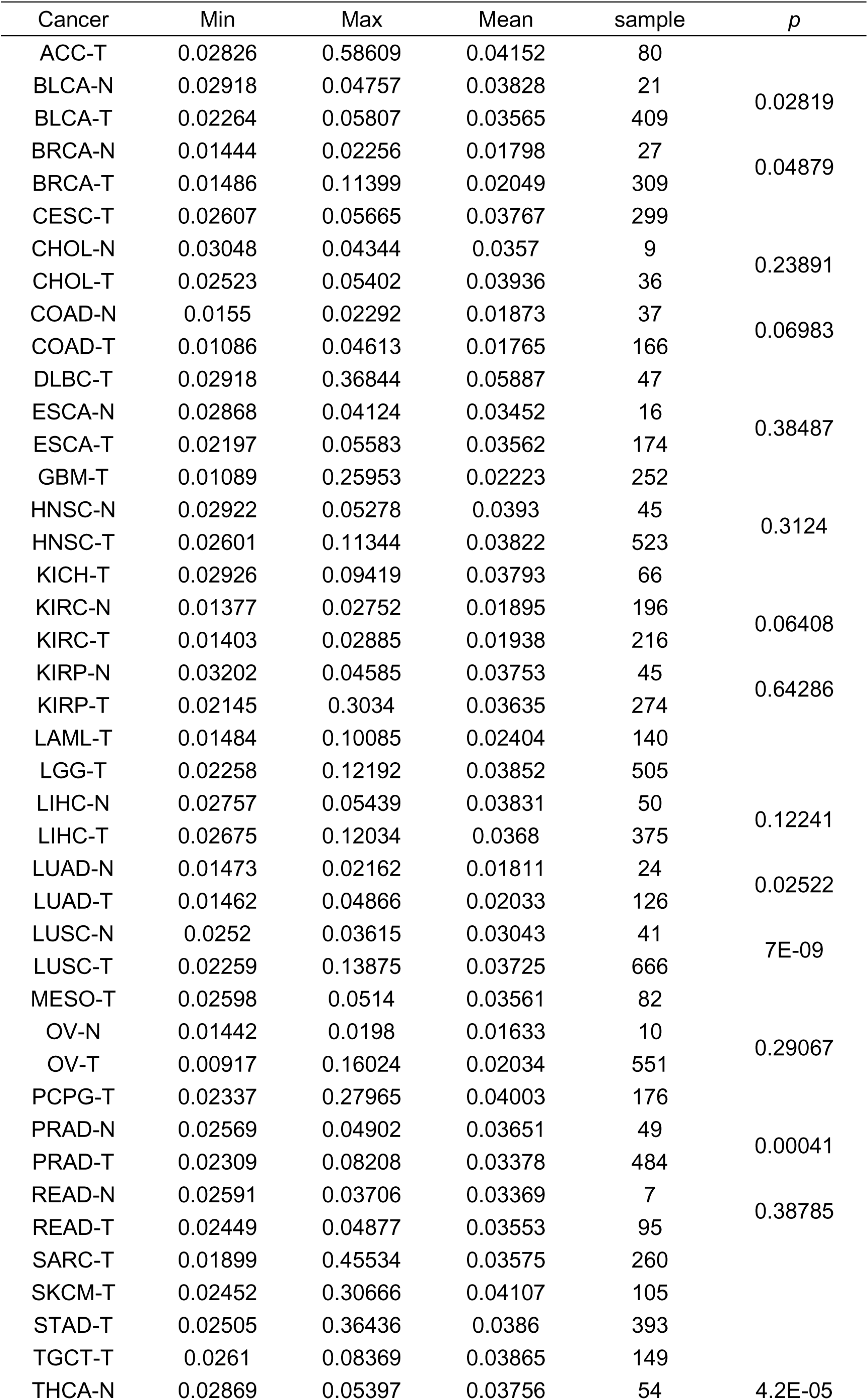
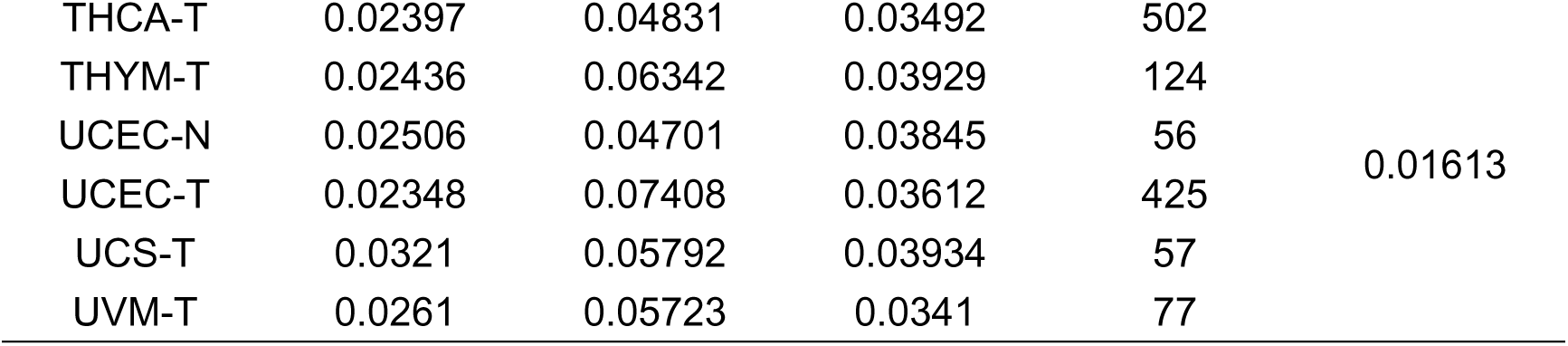
Methylation analysis of TCGA data.

**TABLE IV.** Univariate analysis of OS of LUSC in the TCGA cohort.

| Variables | Category | *p* | 95% CI |  |
| --- | --- | --- | --- | --- |
|  |  |  | Lower limit | Upper limit |
| T | T1 | 0.036 |  |  |
|  | T2 | 0.466 | 0.801 | 1.625 |
|  | T3 | 0.027 | 1.060 | 2.673 |
|  | T4 | 0.031 | 1.069 | 3.866 |
| age |  | 0.073 | 0.999 | 1.033 |
| M |  | 0.007 | 1.147 | 2.424 |
| N | N1 | 0.528 |  |  |
|  | N2 | 0.741 | 0.768 | 1.448 |
|  | N3 | 0.412 | 0.755 | 1.984 |
|  | N4 | 0.202 | 0.613 | 10.129 |
| gender |  | 0.310 | 0.610 | 1.170 |
| stage | stage I | 0.057 |  |  |
|  | stage II | 0.307 | 0.857 | 1.632 |
|  | stage III | 0.030 | 1.040 | 2.155 |
|  | stage IV | 0.056 | 0.973 | 7.279 |
| CDKN1A |  | 0.017 | 1.042 | 1.513 |

